# Protein Synthesis Profiling Enables In Vivo Capture of Ribosome-Associated mRNAs in Yeast

**DOI:** 10.1101/2025.11.17.688896

**Authors:** Cesar Arcasi Matta, Zhi Qi Ten, Simpson Joseph

## Abstract

Translation is a central control point of gene expression, linking nucleotide sequences to functional proteins. Dysregulated translation contributes to diverse diseases, underscoring the need for methods that can directly reveal which transcripts are actively translated. Ribosome profiling, the current gold standard, provides nucleotide-resolution maps of ribosome occupancy but requires laborious purification and sacrifices information on mRNA isoforms and mRNA modifications by restricting analysis to short ribosome-protected fragments.

Here, we introduce Protein Synthesis Profiling (PSP), a proximity-labeling strategy for the transcriptome-wide identification of actively translated mRNAs without the need for ribosome isolation. PSP exploits a fusion of the enzyme APEX2 with the elongation factor eEF2, which transiently associates with ribosomes during elongation, to catalyze selective tagging of mRNAs engaged in translation.

When applied to *Saccharomyces cerevisiae*, PSP captures condition-specific translational programs, recapitulates known stress responses, and expands the detectable repertoire of regulated genes beyond that of existing methods. By preserving full-length transcript features, PSP is scalable, isoform-aware, and broadly adaptable, providing a versatile platform to dissect translational regulation in health and disease.

## Introduction

Dysregulation of mRNA translation contributes to a wide spectrum of human diseases and metabolic syndromes [1,2]. Despite its importance, a comprehensive understanding of which mRNAs are actively translated in healthy versus diseased states remains incomplete. Ribosome profiling is the gold-standard method for measuring translation in vivo, offering nucleotide-resolution maps of ribosome occupancy [3,4]. Ribosome profiling begins by halting mRNA translation, treating the mRNA-ribosome complex with nuclease to generate ribosome footprints, and then recovering the ribosome by ultracentrifugation [4]. Following ultracentrifugation, the ∼30-nucleotide ribosome-protected fragments are purified and put through library preparation for deep sequencing [4]. Although ribosome profiling is commonly used, it has several limitations and challenges. First, it is a laborious technique requiring large material quantities and sophisticated instruments [4]. Second, preparing sequencing libraries from the ∼30-nt ribosome-protected fragments may introduce experimental bias and contaminating footprints, which contribute to mapping ambiguous reads [5]. Third, there is a loss of information regarding mRNA isoforms and post-transcriptional modifications because the short ribosome-protected fragments are not compatible with long-read sequencing methods.

Proximity labeling is a robust method to identify proteins and RNAs localized near a protein of interest within living cells [6]. This technique utilizes engineered enzymes fused to query proteins to generate reactive species that covalently label nearby biomolecules in vivo. Among these enzymes, APEX2, an optimized ascorbate peroxidase, efficiently converts biotin-tyramide into reactive radicals with a short half-life (<1 ms), tagging biomolecules within ∼25 nm [7–13]. Importantly, APEX-seq has demonstrated that APEX2 fusion proteins can effectively label and identify RNAs in living cells through streptavidin enrichment and deep sequencing [11,12,14,15]. Leveraging APEX-seq, we established Protein Synthesis Profiling (PSP), a novel chemical tagging approach that enables transcriptome-wide identification of actively translated full-length mRNAs without the need for fragmentation. PSP utilizes a fusion of ascorbate peroxidase 2 (APEX2) with eukaryotic elongation factor 2 (eEF2), which transiently interacts with ribosomes during the elongation cycle of protein synthesis. Upon activation, APEX2-eEF2 tags nearby molecules, selectively labeling mRNAs engaged in active translation. As a proof of concept, we established PSP in *Saccharomyces cerevisiae*, where it effectively enriched for condition-specific translated transcripts.

## Results

### Rational design of the APEX2-eEF2 fusion for ribosome-proximal labeling

We engineered the APEX2-eEF2 fusion protein by analyzing the structural organization of eEF2 and its interaction with the 80S ribosome during mRNA-tRNA translocation. To identify the optimal fusion site, we examined the high-resolution X-ray crystal structure of eEF2 bound to the eukaryotic 80S ribosome from *S. cerevisiae* (Figure 1A) [16]. APEX2 was fused to the N-terminus of eEF2, positioned away from the decoding center and the mRNA-tRNA complex to avoid disrupting translation. This site is advantageous because domains I and II of eEF2 maintain a stable spatial association with the ribosome, whereas domains III-V undergo conformational changes necessary for translocation [16]. To further minimize interference with GTP hydrolysis or translocation, a flexible 31-amino acid linker was inserted between APEX2 and eEF2. Structural modeling with AlphaFold [17] indicated that ribosome-bound mRNAs remain within a ∼25 nm radius of the APEX2-eEF2 fusion, enabling efficient labeling.

**Figure 1.**
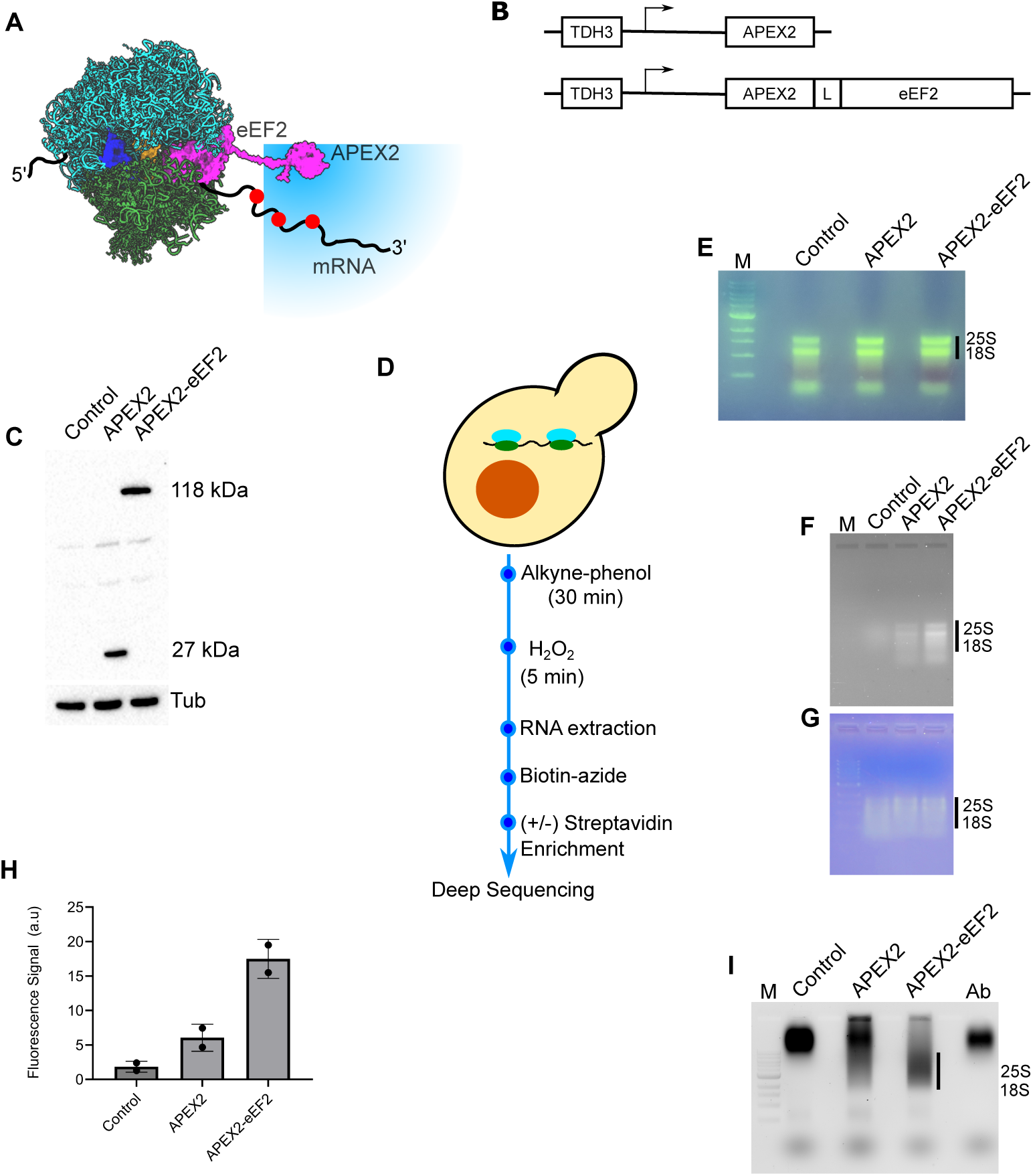
Design and validation of APEX2-eEF2 for RNA tagging in yeast. (A) Structural model of APEX2-eEF2 bound to the translating ribosome. The large ribosomal subunit is shown in cyan, the small subunit in green, and the APEX2-eEF2 fusion protein in magenta. mRNA is shown as a black line, with red circles indicating alkyne-labeled nucleotides generated by proximity labeling. The blue sphere denotes the approximate APEX2 labeling radius (∼20 - 25 nm), illustrating that the APEX2 domain positioned on eEF2 is spatially compatible with efficient labeling of ribosome-associated mRNAs. (B) Schematic of the expression constructs used for cytosolic APEX2 and ribosome-associated APEX2-eEF2, both driven by the constitutive *TDH3* promoter. In the fusion construct, APEX2 is linked to the N-terminus of eEF2 through a flexible linker (L). (C) Western blot validation of APEX2 and APEX2-eEF2 expression in yeast using anti-FLAG detection. Lanes show control (empty vector), APEX2 (∼27 kDa), and APEX2-eEF2 (∼118 kDa). Tubulin (Tub) was used as a loading control. (D) Schematic of the RNA proximity-labeling workflow. Yeast cells were incubated with alkyne-phenol for 30 min, followed by H₂O₂ treatment for 5 min to initiate APEX2-mediated labeling. After quenching, total RNA was extracted, conjugated to biotin-azide by click chemistry, enriched using streptavidin beads when indicated, and used to prepare Illumina sequencing libraries from both total and enriched RNA fractions. (E) Agarose gel analysis of total RNA isolated from control, APEX2, and APEX2-eEF2 cells. Intact 25S and 18S rRNA bands indicate high RNA quality and comparable RNA integrity across samples. M, molecular weight ladder. (F) Detection of alkyne-labeled RNAs following conjugation with fluorescein-azide. Total RNA from control, APEX2, and APEX2-eEF2 cells was subjected to click chemistry and analyzed by agarose gel electrophoresis. Fluorescence was visualized using a Typhoon imager, showing increased RNA labeling in APEX2-eEF2 samples. (G) The same gel shown in (F), stained with SafeStain to confirm equal RNA loading across samples. (H) Quantification of fluorescein-labeled RNA signal from panel (F). Bars represent mean fluorescence intensity ± SD from two independent biological replicates, showing enhanced labeling efficiency for APEX2-eEF2 relative to cytosolic APEX2 and control cells. (I) Gel-shift assay confirming biotin labeling of RNA. RNAs from control, APEX2, and APEX2-eEF2 cells were conjugated to biotin-azide, incubated with anti-biotin-AF488 antibody, and resolved by agarose gel electrophoresis. Antibody-bound biotinylated RNA complexes are indicated by the black bar. Lanes: M, molecular weight ladder; control, RNA from empty-vector cells; APEX2, RNA from APEX2-expressing cells; APEX2-eEF2, RNA from APEX2-eEF2-expressing cells; Ab, antibody only.

For in vivo implementation, the APEX2-eEF2 fusion gene was placed under the control of the strong, constitutive *TDH3* promoter (encoding glyceraldehyde-3-phosphate dehydrogenase) (Figure 1B) and integrated into the chromosome of *S. cerevisiae* strain ZY10 [18]. In parallel, a control strain was constructed in which APEX2 alone was expressed from the same *TDH3* promoter. This TDH3-APEX2 strain served as a background control to distinguish nonspecific RNA labeling by APEX2 from ribosome-associated tagging.

### Enhanced ribosome-proximal RNA labeling with APEX2-eEF2

We examined the expression of APEX2 and the APEX2-eEF2 fusion protein by Western blotting (Figure 1C). A FLAG epitope was present at the N-terminus of APEX2, allowing detection with an anti-FLAG antibody. As expected, the strain expressing APEX2 displayed a band at ∼27 kDa, while the strain expressing the APEX2-eEF2 fusion protein showed a band at ∼118 kDa, consistent with the predicted molecular weights of the respective proteins. No FLAG-reactive band was detected in the control strain lacking APEX2 or APEX2-eEF2 expression. Together, these results confirm the successful in vivo expression of both APEX2 and the APEX2-eEF2 fusion protein.

A previous study demonstrated that alkyne-phenol penetrates the yeast cell wall more efficiently than biotin-phenol, resulting in improved APEX2 labeling [19,20]. To perform proximity RNA labeling, control, APEX2, and APEX2-eEF2 yeast strains were grown to mid-log phase, treated with alkyne-phenol for 30 minutes, and then exposed to hydrogen peroxide for 5 minutes to initiate labeling (Figure 1D). Total RNA was purified and analyzed by agarose gel electrophoresis, which showed intact, high-quality 25S and 18S rRNAs (Figure 1E).

To determine whether the purified RNAs were tagged with alkyne groups by APEX2, we performed copper-catalyzed alkyne-azide click chemistry using a 6-fluorescein-azide (FAM-azide) dye on aliquots of total RNA from each strain. The FAM-labeled RNAs were purified, normalized for concentration, resolved on an agarose gel, and imaged on a Typhoon scanner (Figure 1F). Both APEX2 and APEX2-eEF2 samples showed fluorescent RNA bands compared to the negative control, with the APEX2-eEF2 strain exhibiting ∼3-fold higher fluorescence intensity than APEX2 alone (Figure 1F and 1H). The major fluorescent bands corresponded to 25S and 18S rRNAs, with a diffuse smear below the 18S rRNA consistent with labeled mRNAs. Parallel RNA staining with a non-specific dye confirmed equal RNA loading across samples (Figure 1G). Together, these results indicate that APEX2 non-specifically tags a low fraction of total cellular RNAs (∼5%), whereas APEX2-eEF2 achieves enhanced RNA labeling (∼15%), likely due to its association with translating ribosomes.

We next tested whether alkyne-tagged RNAs could be conjugated to biotin. Total RNA from control, APEX2, and APEX2-eEF2 strains was reacted with biotin-picolyl-azide via click chemistry, followed by RNA purification. The biotinylated RNAs were then incubated with an anti-biotin antibody conjugated to Alexa Fluor 488. The RNA-antibody mixtures were separated by agarose gel electrophoresis and visualized on a Typhoon imager (Figure 1I).

The antibody alone migrated above the 10 kb marker, and a similar band was detected with control RNA. In the APEX2 sample, we primarily observed this free antibody band along with a faint smear extending toward the 25S and 18S rRNAs (near the 3 kb and 2 kb markers). In contrast, the APEX2-eEF2 RNA sample displayed a markedly stronger fluorescent signal corresponding to the rRNAs. This result parallels the direct RNA labeling experiment with FAM-azide, where APEX2-eEF2 showed enhanced fluorescence compared to APEX2. Together, these findings demonstrate that APEX2-eEF2 preferentially tags ribosome-associated RNAs above the background labeling observed with APEX2 alone, further supporting its functional utility for selective RNA labeling on the ribosome.

### Transcriptional and translational reprogramming under amino acid starvation

To validate that the APEX2-eEF2 fusion protein proximity labels ribosomes and potentially ribosome-bound mRNAs, we performed amino acid starvation experiments in yeast. The transcriptional and translational responses to acute amino acid deprivation are well characterized in *S. cerevisiae* and therefore provide a robust benchmark [3,21]. Yeast expressing APEX2 alone or APEX2-eEF2 were grown to mid-log phase and split into two conditions: continued growth in complete medium (control) or transfer to amino acid-deficient medium for 20 min to induce starvation. Following treatment, all cultures were incubated with alkyne-phenol and hydrogen peroxide to initiate APEX2-catalyzed proximity labeling under identical conditions.

Total RNA was isolated from each sample and subjected to copper-catalyzed azide-alkyne cycloaddition with biotin-azide. Biotinylated RNA was then affinity-purified using streptavidin beads, enriching for transcripts labeled non-specifically by APEX2 and, in the case of APEX2-eEF2, those preferentially tagged due to ribosome association. Illumina sequencing libraries were generated from both input (total RNA before enrichment) and enriched fractions for all conditions and constructs. In parallel, a reference library was generated from total RNA isolated from yeast grown to mid-log phase without any treatment, providing a baseline transcriptome.

For each condition, at least three independent biological replicates were generated and used for Illumina library preparation and deep sequencing. Sequencing quality was first assessed using FastQC, followed by alignment of paired-end reads to the *S. cerevisiae* reference genome with STAR [22] and subsequent sorting and indexing with Samtools [23]. On average, each sample yielded approximately 30 to 80 million paired-end reads, with more than 70% mapping uniquely to the yeast genome. Gene-level read counts were quantified using FeatureCounts [24].

To further assess sample quality and reproducibility, count data were normalized using the variance-stabilizing transformation (VST) implemented in DESeq2 [25] (Supplementary Figures 1A and 1B). Principal component analysis (PCA) of the VST-transformed data demonstrated strong concordance among biological replicates, minimal within-group variability, and no extreme outliers across experimental conditions or sequencing batches (Supplementary Figures 1C and 1D). PCA also enabled evaluation of major sources of biological and technical variation, including construct-specific effects (APEX2 versus APEX2-eEF2), nutrient conditions (amino acid-rich versus amino acid-deprived media), and differences between total and ribosome-associated RNA populations.

As expected, the dominant source of variance separated amino acid-starved samples from nutrient-replete controls, confirming that amino acid deprivation induces broad transcriptome remodeling. A clear separation between APEX2 and APEX2-eEF2 samples was also observed, consistent with construct-dependent differences in RNA capture efficiency. Based on these quality-control analyses, three highly concordant biological replicates from each condition were selected for downstream differential expression analysis.

### Unfused APEX2 labels RNA non-selectively, whereas APEX2-eEF2 confers ribosome specificity

Differential gene expression (DGE) analysis was performed using DESeq2 to systematically characterize both transcriptional changes (total RNA) and ribosome-associated signals inferred from enriched RNA fractions in response to amino acid starvation [25]. To first establish the baseline specificity of the labeling chemistry, we evaluated whether unfused APEX2 exhibits any intrinsic bias toward particular RNA populations. Specifically, we compared normalized log₂ fold changes between streptavidin-enriched RNA and corresponding total RNA samples derived from APEX2-expressing cells under both nutrient-replete and amino acid-starved conditions. This analysis directly tests whether APEX2 alone preferentially labels any subset of transcripts independent of ribosome targeting.

Across both conditions, only a small number of genes met statistical thresholds for differential enrichment in the streptavidin-purified fraction relative to total RNA (Figure 2A and 2B). These transcripts were low in number, not reproducibly enriched across conditions, and did not cluster into coherent functional categories by gene ontology analysis. Critically, none of the differentially enriched transcripts corresponded to canonical components of the amino acid starvation response, such as genes involved in amino acid biosynthesis, transport, or general control pathways. Moreover, there was no detectable global shift in enrichment patterns between control and starvation conditions (Figure 2E), indicating that APEX2 labeling does not capture condition-dependent translational remodeling in the absence of ribosome targeting.

**Figure 2.**
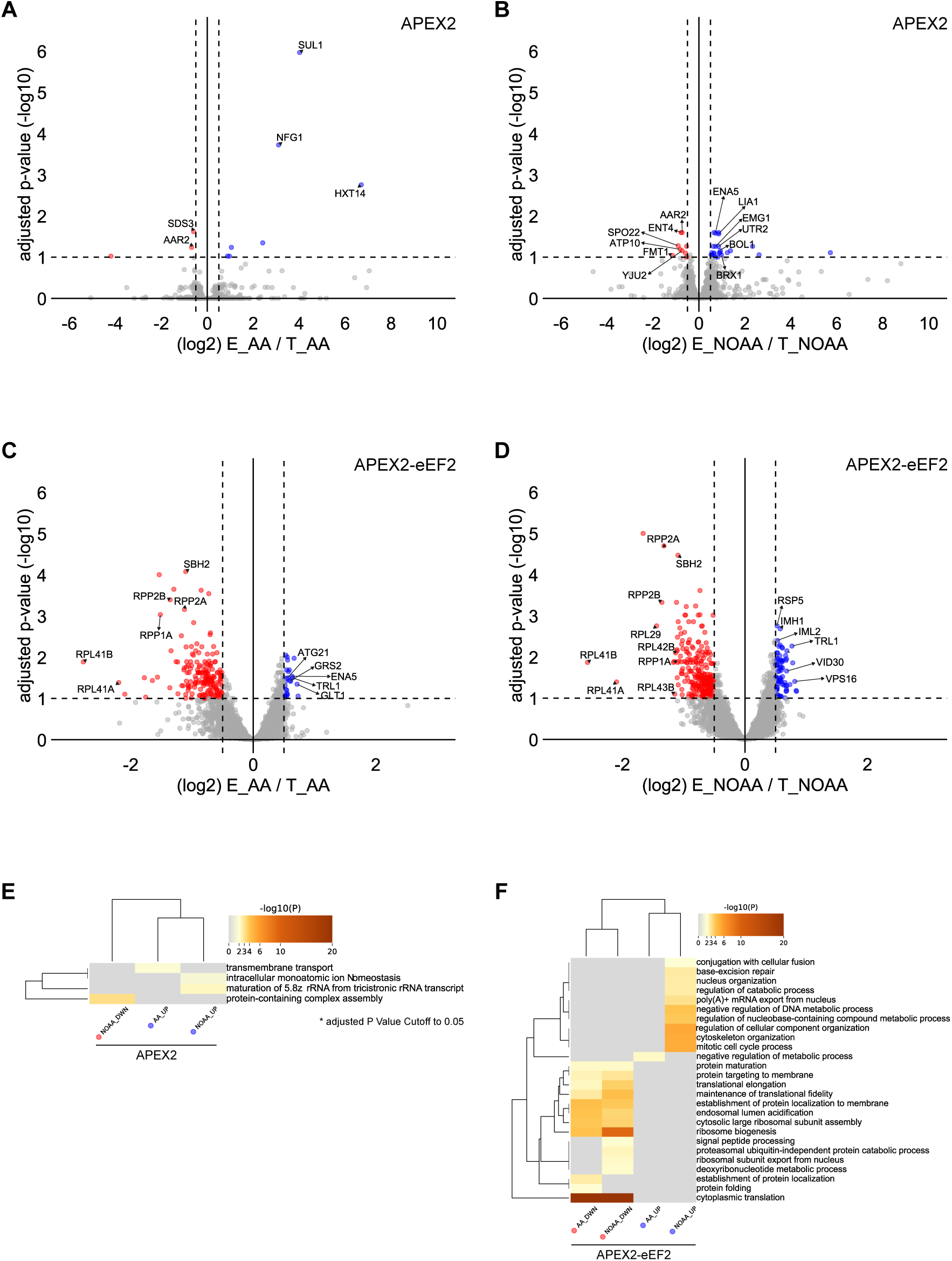
Translational response to amino acid starvation. (A) Volcano plot showing differential enrichment of translated versus total mRNA in amino acid-replete cells (AA) expressing APEX2. (B) Volcano plot showing differential enrichment of translated versus total mRNA in amino acid-starved cells (NOAA) expressing APEX2. (C) Volcano plot showing differential enrichment of translated versus total mRNA in amino acid-replete cells (AA) expressing APEX2-eEF2. (D) Volcano plot showing differential enrichment of translated versus total mRNA in amino acid-starved cells (NOAA) expressing APEX2-eEF2. In panels A - D, each point represents a transcript plotted by log₂ fold change (enriched RNA/total RNA) and adjusted *P* value (-log₁₀ scale). Positive log₂FC values indicate preferential enrichment in the translated RNA fraction (blue), whereas negative values indicate relative depletion (red). Transcripts passing significance thresholds are highlighted, and representative genes are labeled. Notably, APEX2-eEF2 samples show strong depletion of ribosome biogenesis and translation-associated transcripts and enrichment of stress-responsive transcripts under amino acid starvation. (E) Heat map showing Gene Ontology (GO) enrichment analysis of major regulatory classes identified as translationally downregulated (DWN; red) or translationally upregulated (UP; blue) under amino acid-replete (AA) or amino acid-starved (NOAA) conditions. Rows represent significantly enriched biological processes and columns represent the four datasets (AA DWN, NOAA DWN, AA UP, NOAA UP). Color intensity corresponds to enrichment significance (-log₁₀ *P* value), with stronger enrichment shown in darker shades. Prominent repressed categories include cytoplasmic translation, translational elongation, ribosome biogenesis, and maintenance of translational fidelity

Taken together, the near absence of biologically meaningful differentially enriched transcripts demonstrates that unfused APEX2 labels cellular RNAs in a largely non-selective manner. This behavior is consistent with diffuse cytoplasmic proximity labeling driven by enzyme diffusion rather than spatial confinement to translating ribosomes. These results establish a critical negative control, confirming that any enrichment patterns observed with the APEX2-eEF2 fusion arise from ribosome-associated proximity labeling rather than intrinsic sequence or structural biases of the APEX2 chemistry.

In contrast to the non-selective labeling observed with unfused APEX2, analysis of APEX2-eEF2-expressing cells revealed extensive differential enrichment between streptavidin-enriched and total RNA fractions under both nutrient-replete and amino acid-starved conditions (Figure 2C and 2D). Using the same DESeq2 framework described above, we compared normalized log₂ fold changes (enriched versus total RNA) to quantify transcript-specific enrichment associated with the fusion protein. In both conditions, many transcripts passed statistical thresholds for differential enrichment, indicating that tethering APEX2 to eEF2 confers substantial selectivity in RNA labeling.

Functional annotation of these differentially enriched transcripts revealed a striking and biologically coherent partitioning (Figure 2F). Transcripts depleted from the enriched fraction were strongly associated with ribosome biogenesis, rRNA processing, and core components of the cytoplasmic translation machinery. These classes of mRNAs are well known to be translationally repressed during stress and are therefore expected to exhibit reduced ribosome occupancy. In contrast, transcripts enriched in the streptavidin fraction were predominantly linked to amino acid biosynthesis and transport, intermediary metabolism, and other stress-adaptive pathways that are transcriptionally and translationally induced during amino acid deprivation. Gene ontology analysis confirmed that these enriched categories align closely with canonical starvation-responsive programs, in sharp contrast to the absence of such signatures in APEX2-only controls [26].

This reciprocal pattern of depletion and enrichment is consistent with selective proximity labeling of ribosome-associated mRNAs mediated by the APEX2-eEF2 fusion. Given that eEF2 engages elongating ribosomes during translocation, the observed enrichment profile reflects the distribution of actively translated transcripts rather than total transcript abundance. Importantly, the magnitude and coherence of these shifts across biological replicates indicate that the signal is not driven by stochastic labeling or transcript abundance biases but instead captures condition-dependent changes in translational engagement.

Notably, even under nominally nutrient-replete conditions, we detected a modest but reproducible enrichment of stress-responsive transcripts in the APEX2-eEF2 dataset. This observation suggests that the alkyne-phenol and hydrogen peroxide treatment used to initiate proximity labeling may introduce a mild metabolic perturbation or transiently alter translational homeostasis. However, this effect is limited in scope and does not obscure the dominant condition-specific signatures observed upon amino acid starvation. Rather, it highlights the sensitivity of the approach to subtle shifts in translational state, further supporting the conclusion that APEX2-eEF2 reports on ribosome-associated mRNA populations in vivo.

### Interaction modeling isolates ribosome-associated transcripts from non-specific APEX2 labeling

To distinguish ribosome-dependent labeling from residual non-specific signal, we implemented a DESeq2 interaction model that jointly accounts for construct identity (APEX2 versus APEX2-eEF2) and RNA fraction (streptavidin-enriched versus total RNA). This framework tests whether enrichment of a given transcript in the streptavidin fraction is specifically dependent on the eEF2 fusion, rather than reflecting baseline labeling observed with APEX2 alone. By modeling these variables simultaneously, DESeq2 corrects for library size, sequencing depth, and gene-specific dispersion while isolating the component of enrichment uniquely attributable to ribosome tethering. Importantly, this approach does not rely on post hoc subtraction of background.

Instead, it partitions variance at the model level, filtering out transcripts that are equivalently labeled in APEX2 and APEX2-eEF2 samples, and retaining those that show a construct-dependent enrichment shift. As a result, diffuse cytoplasmic labeling contributes minimally to the final signal, yielding a more stringent and interpretable estimate of ribosome-associated RNA.

Application of this interaction model produced a refined and biologically coherent set of differentially enriched transcripts (Figure 3A and 3B). Consistent with the analyses above, transcripts depleted from the enriched fraction were strongly associated with ribosome biogenesis, rRNA processing, and core components of the cytoplasmic translation machinery, consistent with reduced ribosome engagement during stress. In contrast, transcripts exhibiting preferential enrichment in APEX2-eEF2 samples were predominantly linked to amino acid biosynthesis and transport, intermediary metabolism, and other starvation-responsive pathways. Gene ontology analysis [26] confirmed strong concordance with canonical amino acid starvation programs (Figure 3C and Supplementary Figure 2). Together, these results demonstrate that the interaction-based framework effectively normalizes for background labeling and selectively resolves ribosome-associated mRNAs, providing a robust statistical foundation for PSP-based measurements of translational regulation in vivo.

**Figure 3.**
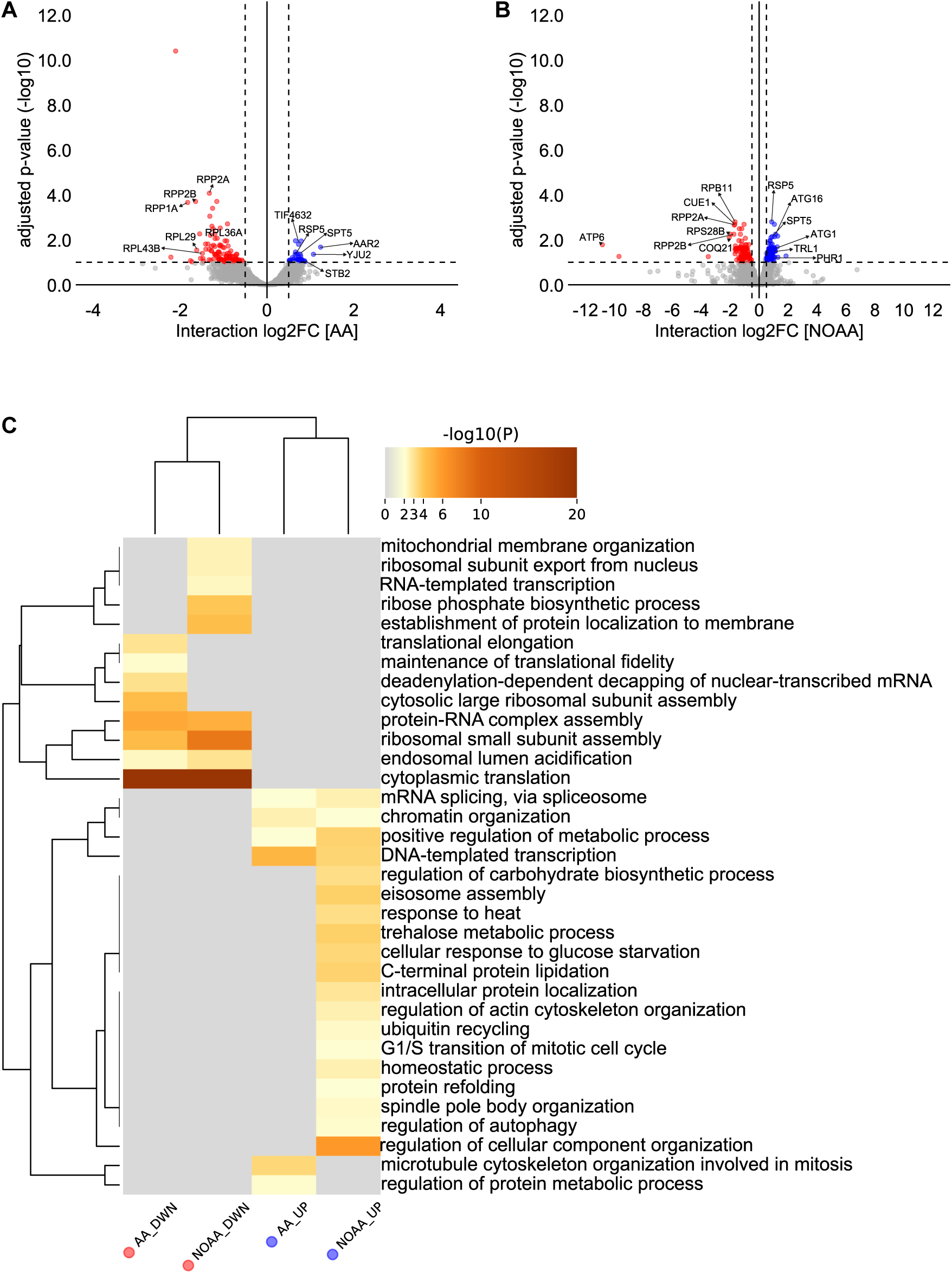
Differential translational regulation revealed by interaction modeling during amino acid starvation. (A) Volcano plot showing transcripts with significant translational regulation in amino acid–replete cells (AA) identified using DESeq2 interaction modeling. The x-axis represents the interaction term log₂ fold change (interaction log₂FC), which distinguishes translational regulation from transcriptional effects, and the y-axis shows adjusted *P* values (-log₁₀ scale). Positive interaction values indicate preferential translational upregulation, whereas negative values indicate translational downregulation. Representative genes are labeled, with ribosome biogenesis and translation-associated transcripts prominently depleted. (B) Volcano plot showing transcripts with significant translational regulation in amino acid-starved cells (NOAA) identified using DESeq2 interaction modeling. Positive interaction log₂FC values indicate transcripts preferentially enriched in the translated RNA fraction during starvation, including stress-responsive and autophagy-related genes, whereas negative values indicate reduced translation of ribosomal and biosynthetic transcripts. (C) Heat map showing Gene Ontology (GO) enrichment analysis of major regulatory classes identified as translationally downregulated (DWN; red) or translationally upregulated (UP; blue) under amino acid–replete (AA) or amino acid-starved (NOAA) conditions. Rows represent significantly enriched biological processes and columns correspond to the four regulatory classes (AA DWN, NOAA DWN, AA UP, NOAA UP). Color intensity indicates enrichment significance (-log₁₀ *P* value), with darker shades representing stronger enrichment. Translationally repressed categories are enriched for cytoplasmic translation, ribosomal subunit assembly, translational elongation, and maintenance of translational fidelity, whereas induced categories are associated with stress adaptation, autophagy, trehalose metabolism, and cellular response to glucose starvation.

### Anota2seq resolves gene-specific translational regulation beyond DESeq2-based enrichment

Although DESeq2 provides a robust framework for differential expression analysis, including the interaction-based normalization described above, it is not explicitly designed to model translational regulation. Approaches that estimate translation efficiency using simple log₂ ratios of enriched to total RNA are prone to spurious correlations and inflated false positives, especially for low-abundance transcripts where variance is high. While the DESeq2 interaction model effectively removes non-specific labeling and isolates ribosome-associated signal, it does not formally partition transcriptional and translational contributions to gene expression changes. Methods such as anota2seq address this limitation by explicitly modeling the dependence between total and translated RNA, thereby enabling more accurate discrimination between transcriptional regulation, translational regulation, and buffering effects [27–30]. Together, these considerations highlight the need for a complementary statistical framework to resolve gene-specific modes of translational control beyond the global enrichment patterns captured by DESeq2.

To this end, we combined PSP with anota2seq to directly disentangle transcriptional and translational effects at transcriptome scale. We benchmarked this analysis against two landmark studies of amino acid deprivation in *S. cerevisiae*: ribosome profiling by Ingolia et al. [3] and polysome profiling by Smirnova et al. [21]. Anota2seq was applied to jointly model total and ribosome-associated RNA across conditions, leveraging its capacity to account for the inherent covariance between these measurements and to reduce false positives arising from ratio-based approaches [27–30]. To facilitate direct comparison with prior datasets that relied on log₂-ratio metrics, we used nominal *p*-values for rank-based evaluation. Within this framework, genes were classified into three principal regulatory categories: (i) translationally regulated, exhibiting changes in translation efficiency independent of mRNA abundance; (ii) mRNA abundance-driven, dominated by transcriptional changes; and (iii) translationally buffered, in which opposing transcriptional and translational changes stabilize protein output [27–30].

Amino acid starvation induced extensive reprogramming of both transcription and translation. In total, 190 genes were translationally upregulated and 298 translationally downregulated (Figure 4A). Gene Ontology (GO) analysis [26] of the translationally upregulated group revealed enrichment for methionine biosynthesis and mitophagy (Figure 4B). These pathways reflect a nutrient-scavenging program that enhances amino acid uptake, recycling, and de novo synthesis to preserve intracellular amino acid homeostasis through the coordinated activation of autophagy, transport, and anabolic metabolism.

**Figure 4.**
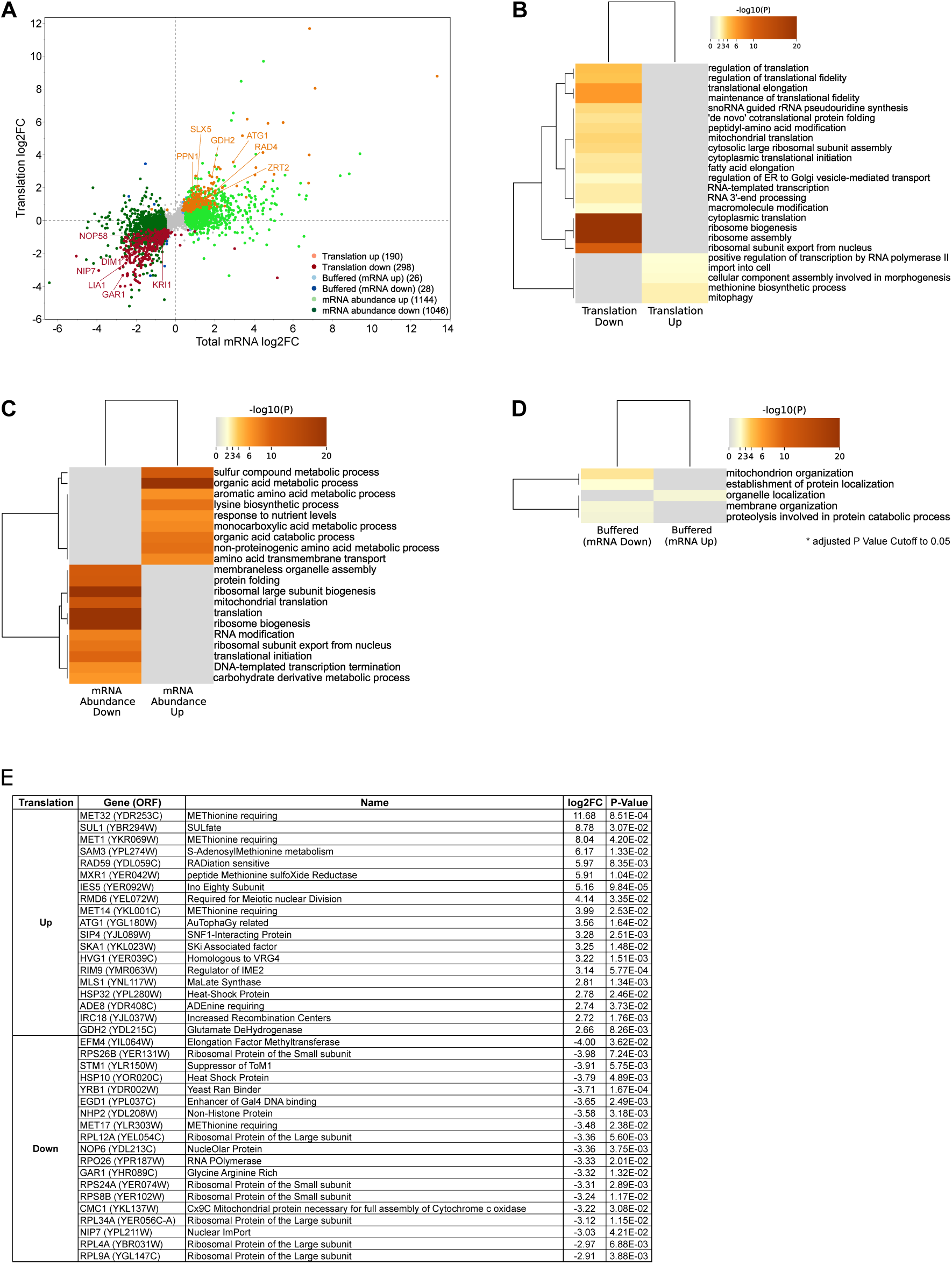
Transcriptional and translational responses to amino acid starvation resolved by anota2seq. (A) Scatter plot comparing changes in total mRNA abundance (x-axis; log₂ fold change) and translation efficiency (y-axis; translational log₂ fold change) between amino acid–replete and amino acid-starved cells. Genes were classified by anota2seq into three major regulatory categories: translational regulation (translation up and translation down), mRNA abundance changes (mRNA up and mRNA down), and buffering (buffered mRNA up and buffered mRNA down), with each class color-coded as indicated. Representative genes are labeled to highlight transcripts showing strong translational activation (e.g., *ATG1*, *GDH2*, *ZRT2*) or repression (e.g., *NIP7*, *GAR1*, *NOP58*). Dashed lines indicate thresholds used for classification. (B) Heat map showing Gene Ontology (GO) enrichment analysis of translationally regulated genes identified by anota2seq, separated into translationally downregulated and translationally upregulated classes. Repressed categories are enriched for cytoplasmic translation, ribosome biogenesis, translational fidelity, and ribosomal subunit assembly, whereas induced genes are associated with methionine biosynthesis and stress adaptation. (C) Heat map showing GO enrichment for genes regulated primarily through changes in mRNA abundance (mRNA abundance down and mRNA abundance up). Downregulated categories are enriched for ribosome biogenesis, mitochondrial translation, and ribosomal subunit export, whereas upregulated categories are associated with sulfur compound metabolism, amino acid biosynthesis, and nutrient-responsive metabolic processes. (D) Heat map showing GO enrichment for buffered genes, in which transcriptional changes are offset at the level of translation (buffered mRNA down and buffered mRNA up). Buffered categories include mitochondrial organization, membrane organization, establishment of protein localization, and proteolysis involved in peptide catabolic processes. (E) Representative examples of significantly affected genes with large changes in translation efficiency identified by anota2seq, together with their open reading frame (ORF), annotated function, log₂ fold change, and adjusted *P* value. Translationally upregulated genes include sulfur assimilation and methionine biosynthesis factors such as *MET32*, *SUL1*, *MET1*, and *MET14*, as well as stress-responsive genes such as *ATG1* and *RAD59*, whereas translationally downregulated genes are enriched for ribosomal proteins and ribosome biogenesis factors, including *RPS26B*, *RPL12A*, *RPL9A*, *NIP7*, and *NOP6*.

In contrast, translationally downregulated genes were strongly enriched for ribosome biogenesis, cytoplasmic translation, ribosomal assembly, and subunit export (Figure 4B). This widespread repression of ribosome-related functions likely represents a resource-conservation strategy that suppresses energetically costly protein synthesis during nutrient limitation.

At the mRNA abundance level, starvation triggered broad changes, with 1,144 genes upregulated and 1,046 downregulated. Upregulated genes were associated with amino acid metabolism, nutrient signaling, and amino acid transport, whereas downregulated genes mirrored translational repression, encompassing ribosome biogenesis, cytoplasmic translation, and mitochondrial translation (Figure 4C and Supplementary Figure 4). These parallel trends underscore the coordinated downshift in global biosynthetic capacity.

Finally, 54 genes displayed translational buffering, including 26 transcriptionally upregulated and 28 transcriptionally downregulated transcripts whose translation levels remained stable. Buffered upregulated genes were enriched for organelle localization, while buffered downregulated genes - linked to mitochondrial organization and proteolysis - may enable rapid translational reactivation upon nutrient recovery (Figure 4D).

### PSP recapitulates and extends known starvation responses

At the gene level, PSP analysis revealed that amino acid starvation selectively enhances translation of MET32, SUL1, MET1, SAM3, MXR1, and MET14, core regulators and enzymes of the methionine and sulfur assimilation network (Figure 4E). Their coordinated induction promotes sulfate import and methionine biosynthesis, complementing the well-characterized Gcn2-Gcn4 transcriptional program [31,32] and uncovering an additional translational layer that sustains homeostasis of sulfur-containing amino acids. PSP also detected increased translation of RAD59, IES5, ATG1, and HSP32, implicating elevated DNA repair capacity, chromatin remodeling, autophagy initiation, and protein quality control as auxiliary stress-mitigation pathways (Figure 4E).

Conversely, translation of EFM4, RPS26B, STM1, RPL12A, NOP6, RPS24A, RPS8B, RPL34A, NIP7, RPL4A, and RPL9A was strongly repressed (Figure 4E), consistent with the conserved energy-saving response that curtails ribosome biogenesis and protein synthesis under nutrient limitation [33–35]. Notably, several of these genes encode ribosomal proteins or assembly factors whose starvation-induced repression was not fully resolved by previous approaches. Together, these findings demonstrate that PSP robustly captures the canonical starvation program while unveiling previously obscured layers of translational regulation that fine-tune metabolic reallocation and proteome remodeling during nutrient stress.

### PSP shows strong global concordance

To evaluate concordance with existing methods, we benchmarked PSP against the Ingolia (ING) [3] and Smirnova (SMR) [21] datasets. ING identified 155 translationally upregulated and 101 downregulated genes, while SMR reported 334 and 240, respectively. A one-way Mann-Whitney U test revealed strong global concordance between PSP and both ING and SMR (Figure 5A and 5B; *P* < 10⁻² - 10⁻⁴). However, a subset of transcripts classified as translationally regulated in ING or SMR showed minimal translation-specific effects in PSP, with approximately 47-58% instead falling into the mRNA abundance class. This redistribution is consistent with anota2seq’s ability to model covariance between total and translated RNA, thereby improving the separation of transcriptional effects from bona fide translation-specific regulation [28,30]. It is also biologically plausible, given prior evidence that severe amino acid starvation in yeast elicits broadly coordinated transcriptome-translatome responses, with more selective translational regulation superimposed on that global program [21,36].

**Figure 5.**
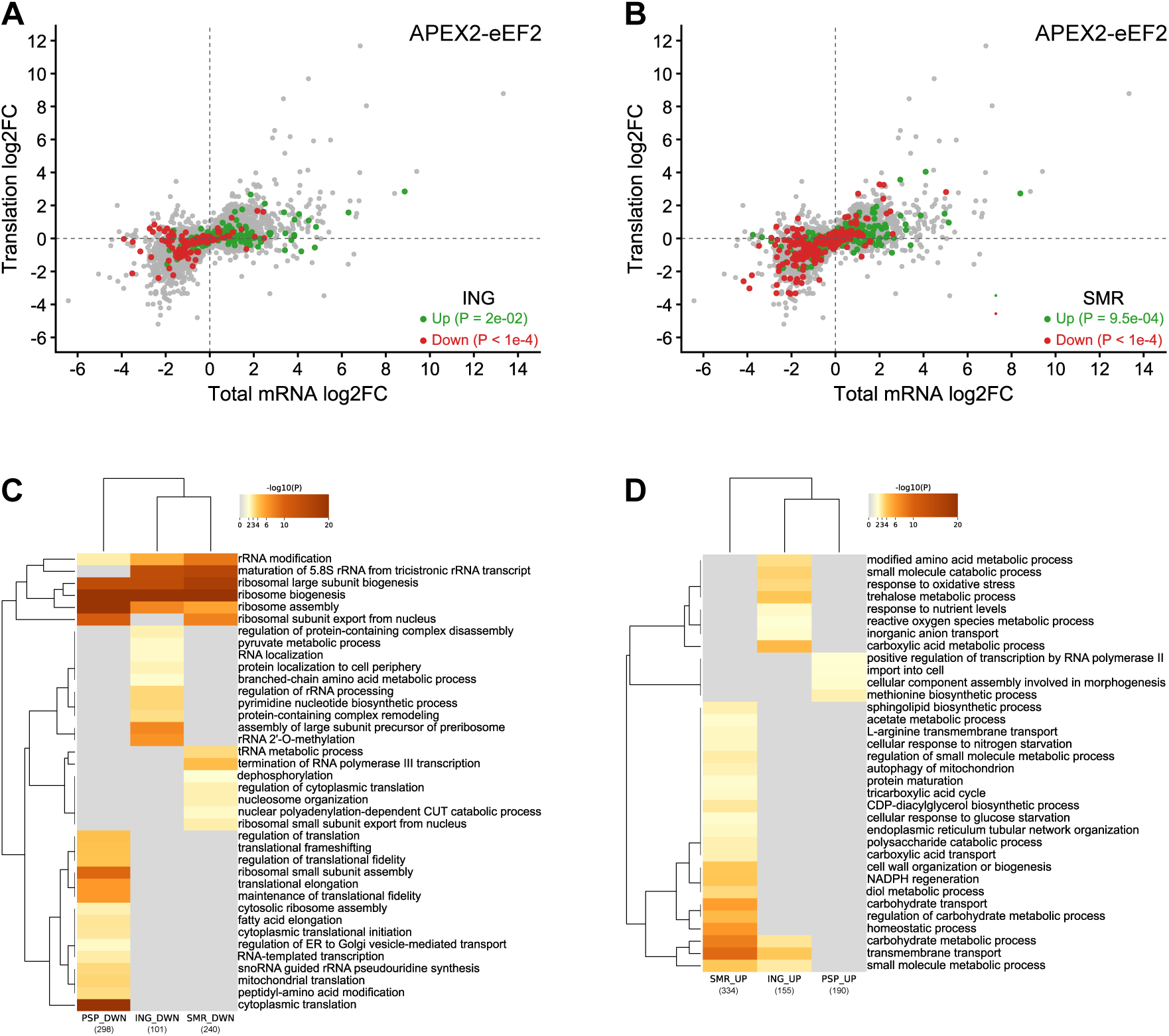
Global concordance among PSP, ribosome profiling, and polysome profiling datasets. (A) Scatter plot comparing PSP-derived changes in total mRNA abundance (x-axis; log₂ fold change) and translation efficiency (y-axis; translational log₂ fold change) with translationally regulated genes identified by ribosome profiling from Ingolia et al. (ING) overlaid. All PSP transcripts are shown in gray as in Figure 4A, whereas ING genes classified as translationally upregulated (green) or downregulated (red) are highlighted. Despite limited gene-level overlap, ING-regulated genes show significant directional concordance with PSP, with translationally upregulated genes enriched in the positive translation quadrant and translationally downregulated genes enriched in the negative translation quadrant. Significance was assessed using a one-sided Mann-Whitney U test (*P* values shown). (B) Scatter plot comparing PSP data with translationally regulated genes identified by polysome profiling from Smirnova et al. (SMR), displayed as in panel A. SMR translationally upregulated genes (green) and downregulated genes (red) similarly show significant directional agreement with PSP, supporting conserved global translational responses across platforms. Significance was assessed using a one-sided Mann-Whitney U test (*P* values shown). (C) Heat map showing Gene Ontology (GO) enrichment analysis of translationally downregulated genes identified by PSP, ING, and SMR. Rows represent significantly enriched biological processes, and columns correspond to the three datasets (PSP, ING, and SMR). Color intensity indicates enrichment significance (-log₁₀ *P* value), with darker shades representing stronger enrichment. Shared repressed categories include ribosome biogenesis, ribosome assembly, rRNA processing, translational elongation, cytoplasmic translation, and maintenance of translational fidelity, demonstrating strong pathway-level concordance despite limited overlap in individual genes. (D) Heat map showing GO enrichment analysis of translationally upregulated genes identified by PSP, ING, and SMR. Rows represent significantly enriched biological processes, and columns correspond to the three datasets. Common induced categories include amino acid metabolic processes, methionine biosynthesis, response to nutrient limitation, trehalose metabolism, autophagy, and broader cellular stress-response pathways, consistent with adaptive translational reprogramming during amino acid starvation. Numbers in parentheses indicate the total number of significantly regulated genes in each class for each dataset.

GO analysis further showed strong concordance in the pathways subject to translational regulation across datasets. Among downregulated gene sets, all three methods showed strong enrichment for ribosome biogenesis and ribosome assembly, consistent with repression of ribosome-related pathways as a conserved response to amino acid starvation (Figure 5C). PSP, however, uniquely resolved additional repressed categories containing genes not detected in ING or SMR, including cytoplasmic translation (for example, SUI2, NIP1, and RPS3), translational fidelity (ASC1, RPS31, and RPL3), and programmed ribosomal frameshifting (HYP2, ANB1, and SSB2). These processes are consistent with active modulation of ribosome function and decoding accuracy during amino acid limitation [33–35,37,38].

Conversely, GO terms uniquely associated with genes upregulated in PSP, but not in ING or SMR, defined a broader adaptive-response program, including sulfur amino acid and methionine biosynthesis (STR3, MET4, MET14), nutrient import and trafficking (MUP1, YPT52, ECM21), transcriptional activation (STP2, DAL81, MET32), mitophagy (ATG1, ATG17, ATG4), and broader cell-remodeling processes (MSO1, SPO21, SPS100) (Figure 5D). Notably, no enriched GO terms were shared among the PSP-, ING-, and SMR-upregulated gene sets. Nevertheless, gene-level analysis of concordantly upregulated genes showed agreement with both reference datasets. The major differences were in the composition of the upregulated signatures: ING reflected a relatively compact program associated with cell wall or spore wall remodeling, membrane metal transport, and amino acid metabolism, whereas SMR reflected a broader stress-adaptive program involving autophagy and organelle remodeling, endosomal trafficking, DNA repair and genome maintenance, and membrane, lipid, and metal homeostasis. PSP captured features of both developmental remodeling and stress-adaptive regulation.

This asymmetry relative to the downregulated sets is consistent with prior yeast amino acid-starvation studies, in which translational repression is repeatedly centered on ribosome biogenesis and protein-synthesis functions, whereas translational induction is more heterogeneous, reflecting early scavenging and metabolic adaptation and later Gcn4-dependent biosynthetic activation [3,21,36]. Overall, PSP-specific enrichments point to an expanded translational control program integrating amino acid biosynthesis, nutrient import, and carbon mobilization, consistent with adaptive metabolic reprogramming associated with amino acid limitation and the Gcn2/GCN4 pathway [30,38–43].

Together, these analyses confirm that PSP faithfully reproduces the conserved starvation-induced programs of translational repression and metabolic remodeling, while extending their resolution to additional regulatory layers. By directly coupling translation-linked mRNA labeling with deep sequencing, PSP delivers a mechanistic and high-fidelity view of how cells remodel translation in response to nutrient stress.

## Discussion

Translation is the decisive checkpoint between RNA and protein, yet it remains the most challenging layer of gene expression to study at scale. Ribosome profiling delivers nucleotide resolution but fragments transcripts and requires labor-intensive purification, while polysome profiling provides only coarse estimates of translational activity [4,44,45]. More recently, editing-based approaches such as Ribo-STAMP have enabled antibody-free detection of ribosome-associated transcripts by fusing APOBEC1 to small ribosomal subunits, creating a cumulative mutational record of ribosome association over extended labeling periods [46,47]. These approaches provide important advances, particularly for single-cell applications, but their temporal resolution is inherently limited by the requirement for prolonged fusion expression and the accumulation of RNA editing events. In addition, because small ribosomal subunits can remain associated with scanning complexes, post-termination complexes, and 3′ UTR regions, these methods often reflect general ribosome association rather than active elongation.

To overcome these limitations, we developed Protein Synthesis Profiling (PSP), a proximity-labeling approach that directly marks ribosome-associated RNAs in vivo. By fusing APEX2 to eEF2, a factor that transiently engages ribosomes during elongation, PSP selectively labels mRNAs in the act of translation while preserving their full length. Because eEF2 functions specifically during mRNA-tRNA translocation, PSP is mechanistically positioned to report productive translation rather than general ribosome occupancy. The acute APEX2 labeling window further allows rapid capture of translational states in response to environmental perturbations, making PSP particularly well-suited for studying transient stress responses and signaling-dependent translational reprogramming. In contrast to footprint-based methods, PSP preserves intact transcripts, making the method compatible with long-read sequencing and enabling analysis of transcript isoforms, untranslated regions, and RNA modifications that are often inaccessible to ribosome footprinting approaches.

Our proof-of-concept in *S. cerevisiae* demonstrates that PSP can detect both global and gene-specific changes in translation under nutrient stress. In response to amino acid starvation, PSP faithfully captured the canonical translational reprogramming response, including downregulation of ribosome biogenesis, cytoplasmic translation, and translation factor mRNAs, coupled with upregulation of amino acid biosynthesis, amino acid transport, sulfur assimilation, autophagy, and metabolic adaptation pathways. These findings align closely with results from ribosome profiling and polysome profiling [3,21], supporting the biological validity of the approach. Importantly, PSP identified a broader spectrum of regulated genes than either traditional method, including additional categories linked to translational fidelity, programmed ribosomal frameshifting, and specialized translational control that were less apparent in prior datasets. We attribute this expanded coverage to deeper sequencing depth, avoidance of ribosome purification biases, and preservation of full-length transcript integrity. This broader detection highlights PSP’s ability to resolve subtle and previously underappreciated translational changes, providing a more complete view of the adaptive gene expression program during nutrient stress.

Equally significant are the methodological advantages of PSP. The workflow is streamlined and scalable, avoiding labor-intensive fractionation, ribosome purification, and RNase digestion steps that often introduce technical variability in ribosome profiling and polysome profiling workflows. This simplicity improves reproducibility and lowers the technical barrier for adoption across laboratories. Because labeling occurs directly in vivo, PSP minimizes perturbation of native translation states during sample preparation and reduces concerns associated with ribosome runoff or stress-induced artifacts during lysis and fractionation. Furthermore, PSP does not rely on prolonged expression of RNA-editing enzymes or transcript recoding [46,47], thereby avoiding the potential introduction of nonsense mutations, frameshifts, or unintended changes in RNA stability that can complicate interpretation in editing-based systems. The method is also highly versatile and adaptable, readily applicable to diverse experimental designs, stress conditions, and biological systems, and can be transferred to mammalian cells without major technical barriers.

An important analytical consideration is the distinction between transcriptional and translational regulation. Traditional translation efficiency calculations based on simple log-ratio approaches are highly susceptible to spurious correlations and inflated false positives, particularly for low-abundance transcripts. By integrating PSP with anota2seq [27–30], we were able to explicitly model the dependence between total RNA and translated RNA, allowing robust separation of transcriptional regulation, translational regulation, and buffering effects. This framework substantially improves interpretability compared to conventional translational efficiency estimates and strengthens confidence in gene-specific translational assignments. The strong pathway-level concordance observed between PSP and prior ribosome profiling datasets, despite limited gene-level overlap, further underscores the importance of rigorous statistical modeling when interpreting translational control at transcriptome scale.

Looking ahead, PSP has the potential to reshape the study of translational control. By directly labeling ribosome-associated mRNAs in vivo, PSP provides a framework that can be extended to dynamic studies of translation in response to environmental stress, signaling pathways, developmental programs, host-pathogen interactions, and disease states. The rapid temporal resolution of APEX2 labeling is particularly advantageous for capturing short-lived translational responses that are difficult to resolve using cumulative editing-based approaches. In parallel, the preservation of full-length RNAs enables integration with long-read sequencing platforms to study isoform-specific translation, alternative polyadenylation, untranslated region usage, and RNA modification-dependent regulation. These features substantially expand the biological questions that can be addressed beyond what is possible with conventional footprint-based methods.

The method is also inherently compatible with emerging single-cell and spatial transcriptomics platforms, offering a path to high-resolution mapping of translational regulation across heterogeneous tissues, developmental gradients, and tumor microenvironments [48–50].

Although the current implementation was developed in bulk yeast cultures, the conceptual framework of acute proximity labeling combined with RNA capture is highly compatible with future adaptation to single-cell workflows. This ability to bridge transcriptional, translational, and spatial layers of gene regulation represents a unique strength of PSP and positions the method for broad impact across molecular and cellular biology.

In summary, PSP expands the experimental toolkit for investigating translational regulation by combining biochemical specificity with methodological simplicity. Its ability to capture actively elongating ribosomes with high temporal precision distinguishes it from methods that primarily measure cumulative ribosome association. Its enhanced transcript coverage, broad adaptability, and compatibility with cutting-edge sequencing technologies position PSP not merely as a complement to ribosome and polysome profiling, but as a next-generation platform for probing translation in complex biological systems. By lowering technical barriers while increasing biological insight, PSP opens new opportunities to study translation with unprecedented depth, breadth, and resolution, poised to have a transformative impact on molecular biology, systems biology, and biomedical research.

## Methods

### Plasmid construction and yeast transformation

The pTDH3-APEX2 plasmid was generated by PCR amplification of the *APEX2* gene from pJH124 (Addgene #102951) and insertion into p406TDH3 (Addgene #15977) by homologous recombination. To create pTDH3-APEX2-eEF2, the yeast *eEF2* gene was PCR-amplified from pTKB612 (a gift from Dr. Jonathan D. Dinman, University of Maryland) and inserted into pTDH3-APEX2 by homologous recombination. All constructs were verified by whole-plasmid sequencing (Plasmidsaurus).

The pTDH3-APEX2 and pTDH3-APEX2-eEF2 plasmids were linearized with *NcoI-HF* and transformed into *S. cerevisiae* strain ZY10 (W303 background: MATa trp1 leu2 ura3 his3 can1 GAL+ psi+) [18] (a gift from Dr. Brian Zid, UCSD) to achieve chromosomal integration. Transformations were carried out using the Frozen-EZ Yeast Transformation II kit (Zymo Research, Cat# T2001) according to the manufacturer’s protocol. Chromosomal integration was confirmed by PCR and Sanger sequencing.

### Protein extraction and Western blotting

A single yeast colony from a Synthetic Complete (SC) (-Ura) agar plate having 2% D-glucose was inoculated into 3 mL of SC (-Ura) medium with 2% D-glucose and grown overnight at 30°C. The culture was diluted to an OD600 of ∼0.1 in 10 mL of fresh SC (-Ura) medium with 2% D-glucose and incubated at 30 °C until the OD600 reached ∼1.0. Cells were harvested by centrifugation at 4,835 × g for 15 minutes at 4 °C, and the pellet was washed twice with 5 mL 1× PBS (pH 7.4). The washed pellet was resuspended in 200 µL of 0.2 M NaOH, incubated at room temperature for 5 minutes, and centrifuged at 12,000 rpm for 1 minute. The pellet was resuspended in 100 µL of 2× SDS loading buffer (24 mM Tris-HCl pH 6.8, 10% glycerol, 0.8% SDS, 5.76 mM β-mercaptoethanol, 0.04% bromophenol blue) and boiled at 95 °C for 5 minutes. Lysates were centrifuged at 3,000 rpm for 10 minutes at 4 °C, and supernatants were collected.

Protein concentration was estimated by A280 measurement using a Spark Multimode Microplate Reader (Tecan, Switzerland). For SDS-PAGE, 2.5 µg of total protein was loaded onto a 10% gel alongside a pre-stained protein ladder (NEB, Cat# P7706). Electrophoresis was performed in two steps: 100 V for 10 minutes, followed by 180 V for 40 minutes. Proteins were transferred onto a PVDF membrane pre-activated in methanol for 30 seconds, rinsed in deionized water (5×), and equilibrated in transfer buffer (25 mM Tris, 192 mM glycine, 20% methanol) for 15 minutes.

Transfer was performed using a Bio-Rad Trans-Blot SD semi-dry system at 20 V for 1 hour. Membrane was blocked in 5% milk for 1 hour at room temperature with gentle shaking, washed 5 times with 1× TBST (20 mM Tris, 150 mM NaCl, 0.1% Tween 20), and incubated overnight at 4 °C with mouse anti-FLAG Tag monoclonal antibody (1:5,000 dilution) (Invitrogen Cat# MA1-91878) in 5% milk. After washing 3 times with 1× TBST, the membrane was incubated with HRP-conjugated anti-mouse IgG secondary antibody (1:10,000 dilution) (Cell Signaling Technology Cat# 7076) for 1 hour at room temperature. After additional washes, chemiluminescence was developed with Amersham ECL substrate and imaged using a Bio-Rad ChemiDoc MP Imaging System.

To check for loading control, the membrane was stripped in stripping buffer (10 mM β-mercaptoethanol, 62.6 mM Tris-HCl pH 6.8, 2% SDS) at 55 °C for 40 minutes. The blot was reprobed with mouse anti-alpha-tubulin monoclonal antibody (1:8000 dilution) (DSHB Cat# 12G10) in 5% milk for 2 hours at room temperature, followed by the HRP-conjugated anti-mouse IgG secondary antibody (1:10,000 dilution) and detected as described above.

### APEX2-catalyzed alkyne-phenol tagging of RNA

Yeast cells were cultured in 60 mL of SC (-Ura) media with 2% D-glucose at 30 °C with vigorous shaking. Cultures were inoculated at an initial OD600 of 0.1 and grown to mid-log phase (OD600 = 0.6–0.7). Cells were harvested, washed once with 1× PBS (pH 7.4), and resuspended in 500 µL of 1× PBS supplemented with 2.5 mM alkyne-phenol (MedChemExpress LLC, Cat# HY-131442). The suspension was incubated at room temperature with constant shaking for 30 minutes.

To initiate APEX2 labeling, 5 µL of 100 mM hydrogen peroxide was added, briefly mixed by gentle vortexing, and incubated at room temperature for 5 minutes. The reaction was quenched by adding 500 µL of quenching buffer (10 mM NaN_3_, 5 mM Trolox, 10 mM sodium ascorbate in 1× PBS), followed by gentle vortexing. Cells were pelleted by centrifugation at 15,294 × g for 5 minutes at 4 °C, and the supernatant was discarded. The pellet was washed twice with 500 µL of 1× PBS and resuspended in FAE solution (98% formamide, 10 mM EDTA, pH 8).

The suspension was heated at 70 °C for 10 minutes, then centrifuged at 21,000 × g for 2 minutes at room temperature. The RNA-containing FAE supernatant was transferred to a fresh DNA/RNA Lo-Bind 1.5 mL microcentrifuge tube, diluted to 78% formamide with water, and purified using the RNA Clean & Concentrator-100 kit (Zymo Research), according to the manufacturer’s instructions.

The concentration of alkyne-labeled RNA was measured with a Spark Multimode Microplate Reader (Tecan, Switzerland), and RNA quality was assessed by 1% TAE agarose gel electrophoresis. DNase treatment was performed using RQ1 RNase-Free DNase (Promega, Cat# M6101) following the manufacturer’s protocol. DNase-treated RNA was further purified using the Monarch RNA Cleanup Kit (New England BioLabs, Cat# T2040). Final RNA concentration was determined using the microplate reader, quality was verified by agarose gel electrophoresis, and samples were stored at −80 °C.

### Click chemistry labeling of RNA

A total of 100 µg of alkyne-labeled RNA was used for the click reaction. The reaction mixture contained 10 mM Tris-HCl (pH 7.5), 2 mM biotin picolyl azide (Click Chemistry Tools, Cat# 1167), 0.1 mM CuSO₄, 2 mM THPTA, and 10 mM sodium ascorbate. Samples were vortexed, briefly centrifuged, and incubated in the dark at room temperature for 30 minutes with constant shaking.

After incubation, RNA was purified using the Monarch RNA Cleanup Kit (New England BioLabs, Cat# T2040) according to the manufacturer’s instructions. The RNA concentration was determined with a Spark Multimode Microplate Reader (Tecan, Switzerland), integrity was assessed by 1% TAE agarose gel electrophoresis, and samples were stored at −80 °C.

### Fluorescein labeling of RNA

A total of 20 µg of alkyne-labeled RNA was subjected to a click reaction with 2 mM 6-fluorescein-azide (Jena Bioscience, Cat# CLK-80105), as described above. From this reaction, 3 µg of fluorescein-labeled RNA was mixed with 1× loading dye and resolved on a 1% TAE agarose gel at 70 V for 40 minutes. Alternatively, 5 µg of fluorescein-labeled RNA was analyzed by 10% urea-PAGE.

In-gel fluorescence was detected using an Amersham Typhoon Biomolecular Imager (Cytiva, MA, USA) with Cy2 configuration (488 nm laser, 525BP20 filter, 50 µm pixel size).

Fluorescence intensity of dye-labeled RNA was quantified using the BandPeak tool in ImageJ. Following fluorescence imaging, the agarose gel was stained in 30 mL of 1× TAE buffer containing 3 µL SYBR Gold nucleic acid gel stain (Invitrogen) for 1 hour with gentle shaking and visualized under UV illumination to assess total RNA.

### Detection of biotinylated RNA

A total of 10 µg of biotinylated RNA in 1× PBS was incubated with 250 ng of anti-biotin monoclonal antibody (BK-1/39) conjugated to Alexa Fluor 488 (ThermoFisher Cat# 53-9895-82) at room temperature for 30 minutes with constant shaking in the dark. After incubation, 1× agarose loading dye was added, and samples were resolved on a 1% TAE agarose gel alongside a 1 kb DNA ladder (New England BioLabs, MA, USA). Electrophoresis was performed at 80 V for 35 minutes.

Fluorescent imaging was carried out on an Amersham Typhoon Biomolecular Imager (Cytiva, MA, USA) using Cy2 configuration. Following imaging, the gel was stained in 1× PBS (pH 7.4) containing DNA SafeStain (Lamda Biotech, Cat# C138) for 1 hour with gentle rocking and visualized under UV illumination.

### Amino acid deprivation and APEX2 labeling of RNA

Yeast cells were cultured in SC (-Ura) media with 2% D-glucose at 30 °C with vigorous shaking. Cultures were inoculated at an initial OD600 of 0.1 and grown to mid-log phase (OD600 = 0.6–0.7). The culture was divided into two equal volumes and centrifuged at 4,500 × g for 8 minutes at 30 °C to pellet the cells, followed by one wash with 1× PBS and re-centrifugation. One pellet was resuspended in complete SC (-Ura) medium containing 2% D-glucose, while the other was resuspended in SC (-Ura) medium lacking amino acids but supplemented with 2% D-glucose. Both cultures were incubated at 30 °C with vigorous shaking for 20 minutes, and final OD600 values were measured and recorded. Cells were then pelleted and resuspended in 500 µL of 1× PBS supplemented with 2.5 mM alkyne-phenol for APEX2-catalyzed RNA tagging, as described above.

### Enrichment of biotinylated RNA using streptavidin beads

Dynabeads MyOne Streptavidin C1 (Invitrogen Cat# 65001) (5 - 7 µg RNA/µL of beads) was equilibrated at room temperature for 20 minutes, washed, and primed for binding according to the manufacturer’s instructions. A total of 100 µg of biotinylated RNA was incubated with the beads for 1 hour at room temperature with gentle rotation. Following incubation, the bead-RNA mixture was placed on a magnetic stand for 3 minutes to allow bead immobilization, and the supernatant was discarded. The beads were then washed again as recommended by the manufacturer.

For elution, beads were resuspended in 50 µL of elution buffer (95% formamide, 10 mM EDTA, pH 8.2, 1.5 mM D-biotin) by gentle flicking until homogeneous. The suspension was heated sequentially at 50 °C for 5 minutes and 90 °C for 5 minutes. Beads were immobilized on a magnetic stand for 3 minutes, and the supernatant containing enriched biotinylated RNA was collected into a fresh tube. The eluted RNA was diluted to <75% formamide with nuclease-free water and purified using the RNA Clean & Concentrator-25 kit (Zymo Research), following the manufacturer’s instructions. The final RNA concentration was quantified using a Tecan Spark multimode plate reader.

### Illumina library preparation and high-throughput sequencing

Between 300 - 1000 ng of total or enriched biotinylated RNA was used for library preparation with the Illumina Stranded mRNA Prep Ligation Kit (Cat# 20040532), following the manufacturer’s instructions. Four biological replicates per condition were processed independently to ensure reproducibility. Final cDNA libraries were quantified with a Qubit 4

Fluorometer (Invitrogen) and assessed for quality and fragment size distribution using an Agilent 4200 TapeStation System (Agilent Technologies) at the Institute of Genomic Medicine (IGM), UC San Diego. Libraries were pooled equimolarly, multiplexed, and sequenced on an Illumina NovaSeq X Plus 10B platform at IGM, generating a minimum of 30 million paired-end reads per sample.

### Sequence Alignment

Base calling and demultiplexing were performed with Illumina BaseSpace™ and the DRAGEN™ Bio-IT Platform. Sequencing quality was assessed with FastQC (v0.11.9), and all libraries passed Illumina quality control thresholds (Q30, yield); no trimming was required.

Reads were aligned to the Saccharomyces cerevisiae S288C reference genome (R64-1-1 assembly) using STAR (v2.7.11) [22] with the Ensembl release 113 reference genome and annotation [51]. Genome indices were generated with STAR’s --runMode genomeGenerate using the reference genome FASTA (Saccharomyces_cerevisiae.R64-1-1.dna_sm.toplevel.fa.gz) and annotation GTF (Saccharomyces_cerevisiae.R64-1-1.113.gtf.gz). The --sjdbGTFfile parameter was used to incorporate splice junctions, and --genomeSAindexNbases 10 was specified for the yeast genome. Alignments were performed using the following parameters: --readFilesCommand zcat, --outSAMtype BAM SortedByCoordinate, --quantMode TranscriptomeSAM, and --outFilterMismatchNmax 2. Coordinate-sorted BAM files were indexed with Samtools (v1.13) [23].

### Read Quantification and Processing

Aligned reads were quantified at the gene level using featureCounts (v2.0.3, Subread) [24] with the S. cerevisiae Ensembl release 113 annotation. Downstream analyses were restricted to protein-coding genes. For every RNA fraction (total and enriched), we combined count tables from various constructs (APEX, APEX2-eEF2, and no-construct control) across different conditions (AA and NOAA) into a single matrix. Column names were standardized, and merged matrices were checked for consistency across samples and conditions. Gene identifiers were subsequently standardized to SGD systematic names for downstream analyses [52].

### Data Quality Control

Preliminary analyses assessed replicate concordance, identified sample outliers, and evaluated potential batch effects to distinguish biological differences between constructs, conditions, and RNA fractions. Count tables were merged by gene and restricted to protein-coding genes defined in the Ensembl release 113 annotations; genes with fewer than 10 total counts across all samples were excluded. Sample metadata (condition, batch, RNA type) were joined to the count matrix for DESeq2 [25]. Counts were normalized with DESeq2 (median-ratio method) and transformed using the variance-stabilizing transformation (VST). VST data were used for quality control analysis, including principal component analysis, sample-to-sample distance heatmaps with hierarchical clustering, and summary assessments of count distribution and dispersion patterns (Supplementary Figure 1).

### Data Preprocessing

Gene annotations were standardized using the Saccharomyces Genome Database (SGD; R64.5.1) [52], and downstream analysis was restricted to verified ORFs (excluding dubious/uncharacterized entries). Processed featureCounts tables correspond to total RNA (T) and translated/enriched RNA (P) from both APEX2 and APEX2-eEF2 constructs, and non-construct control, were subset to the verified ORF set shared across datasets and aligned to a common gene order. Sample metadata (condition: AA vs. NOAA; RNA type: T vs. P) were consistent across matrices, and sample ordering and experimental design were validated before downstream analysis. For anota2seq analysis, datasets were generated from the paired total and enriched count matrices using RNA-seq settings in anota2seq (v1.30.0) [27]. Counts were TMM-log2 normalized, and genes with zero counts across all samples were removed. The differential contrast of interest (AA versus NOAA) was specified, and its direction was verified prior to statistical testing.

### DESeq2 Differential Gene Expression Analysis

Differential expression analysis was performed with DESeq2 using the preprocessed unified count matrix and the corresponding sample metadata. Outlier samples identified during quality-control analyses were removed before model fitting. Following outlier removal, the analysis included the principal experimental groups defined by construct APEX2 or APEX2-eEF2, condition (AA or NOAA), and RNA fraction, encoded as total (TOT) or enriched (ENR), together with baseline control consisting of untreated total RNA (C0_TOT_BL). These samples were encoded as a nine-level composite factor (group). Counts retained after preprocessing and verified-ORF filtering were analyzed in a single DESeq2 model fit across all samples using DESeq, enabling the extraction of all planned pairwise comparisons within a shared normalization and dispersion framework. Differential expression results were obtained using explicit contrasts of the form contrast = c(“group”, numerator, denominator). This approach was used to generate multiple biologically relevant comparison classes, including: (i) construct comparisons (APEX2-eEF2 versus APEX2) within each condition and RNA fraction; (ii) condition comparisons (NOAA versus AA) within each construct and RNA fraction; (iii) enrichment comparisons (ENR versus TOT) within each construct and condition; and (iv) selected comparisons of total RNA samples from APEX2 or APEX2-eEF2 relative to the baseline control group C0_TOT_BL. Positive log2 fold-change values indicate higher abundance in the numerator group, whereas negative values indicate higher abundance in the denominator group. Statistical testing was performed with alpha = 0.1. For downstream reporting and visualization, genes were prioritized using an adjusted P-value threshold of 0.1 and an absolute log2 fold-change threshold of 0.5. Volcano plots were generated from DESeq2 result tables to visualize contrast-specific expression changes. Significance coloring was based on the same thresholds used for reporting (adjusted P ≤ 0.1 and absolute log2 fold change ≥ 0.5): genes with higher abundance in the numerator group were colored blue, genes with higher abundance in the denominator group were colored red, and non-significant genes were shown in gray. For annotation, the top 10 genes in each direction ranked by adjusted P value and the top 10 genes in each direction ranked by absolute log2 fold change were selected for labeling.

### Interaction-model analysis of construct-dependent RNA enrichment

To distinguish construct-dependent RNA enrichment from construct-independent background labeling, DESeq2 interaction analyses were performed separately for each growth condition (AA and NOAA). Within each condition, only samples from constructs APEX2 and APEX2-eEF2 and fractions TOT and ENR were included. Counts were modeled using the design ∼ Construct + SeqType + Construct: SeqType, with C2 and TOT specified as reference levels. The interaction term, therefore, tested whether the magnitude of enrichment in the streptavidin-purified fraction (ENR versus TOT) differed between constructs, corresponding to the contrast (APEX2-eEF2_ENR - APEX2-eEF2_TOT) - (APEX2_ENR – APEX2_TOT). Genes were classified as significant at an adjusted P value ≤ 0.1 and an absolute interaction log2 fold change ≥ 0.5.

### Anota2seq Analysis of Differential Translation

Differential translation analyses were performed with anota2seq v1.30.0 using RNA-seq settings as described in Larsson et al. (2010) [30]. To assess concordance during benchmarking, an initial nominal APV threshold of maxP = 0.05 was applied. Reliability safeguards included joint modeling of paired enriched (ENR) and total (TOT) RNA measurements, random-variance moderation, routine quality-control procedures, and residual-outlier testing (Supplementary Figure 3). A minimum APV effect-size threshold of |log2| ≥ 0.5 was required, and predefined slope bounds were applied for translation and buffering/offsetting analyses. Results were generated for four analysis classes: translation, buffering, translated mRNA abundance, and total mRNA abundance. Regulatory modes, including translation, buffering/offsetting, and mRNA abundance effects, were assigned according to the standard anota2seq classification framework and summarized using diagnostic plots. For the NOAA versus AA contrast, candidate genes were selected using the thresholds described above across all four analysis classes. False discovery rate-adjusted P values (apvRvmPAdj) are reported for each gene. Complete and filtered result tables are provided in the supplementary data (Supplementary Table 1).

### Gene Ontology (GO) Analysis

GO enrichment was performed with Metascape (v3.5) [26] using S. cerevisiae SGD IDs with built-in ID conversion. Single-list analyses were conducted for each regulatory mode and direction (translation, buffering, mRNA abundance; up/down) (Supplementary Tables 2-4). Unless noted otherwise, tests were restricted to Biological Process terms, used the whole-genome background, and used default enrichment analysis settings. For cross-study comparisons involving the present study (PSP), the Ingolia et al. dataset (ING) [3], and the Smirnova et al. dataset (SMR) [21], we used Metascape multi-list enrichment on the categorized translation-mode gene sets, analyzing up-regulated (PSP_Up, ING_Up, SMR_Up) and down-regulated (PSP_Down, ING_Down, SMR_Down) lists separately. Results include per-list and combined-list enrichments with standard term clustering. Single- and multi-list result tables are provided as supplementary data (Supplementary Tables 5 and 6). Genes not handled by Metascape were annotated via SGD.

### Comparative Analysis

We compared PSP results with two reference datasets: the polysome-profiling dataset of Smirnova et al. (SMR) [21] and the ribosome-profiling dataset of Ingolia et al. (ING) [3]. Reported lists of translationally regulated genes were mapped to SGD systematic names and aligned to PSP gene IDs. For directional concordance analysis, we followed the comparison framework used by Ingolia et al. [3]. Translationally up- and down-regulated gene sets from each reference were tested for directional shifts in PSP translation APV log2 effects relative to background genes. The PSP comparison universe included all genes in the anota2seq output with non-missing ORF, total mRNA effect, and translation effect values. ORF identifiers were uppercased, trimmed, and deduplicated to a single entry per gene, and no additional significance filtering was applied before testing. For each reference-direction pair, PSP translation APV log2 effects were compared between genes in the reference set and all remaining genes in the PSP universe using one-sided Mann-Whitney U tests (Up > background; Down < background). We additionally recorded the U statistic, the tie-corrected Z score, the one-sided P value, the area under the curve, the rank-biserial correlation, and the number of reference genes matched in the PSP universe. Scatterplots of PSP total mRNA APV log2 effects versus translation APV log2 effects were generated for each reference, with Up and Down genes overlaid on background genes.

### Quantification and Statistical Analysis

Analyses were performed in R using DESeq2 and anota2seq. GO enrichment analyses were performed using Metascape. Statistical testing included Wald tests within DESeq2 models, false discovery rate adjustment for multiple testing, one-sided Mann-Whitney U tests for directional concordance analyses, and hierarchical clustering for sample-level quality-control analyses.

## ACKNOWLEDGMENTS

This work was supported by the National Institutes of Health (R35GM141864 to S.J.). This publication includes data generated at the UC San Diego IGM Genomics Center utilizing an Illumina NovaSeq X Plus that was purchased with funding from a National Institutes of Health SIG grant (#S10 OD026929).

## Contributions

C.A.M., Z.Q.T., and S.J. designed the experiments. C.A.M. and Z.Q.T. performed the experiments, and all authors discussed the results. C.A.M., Z.Q.T., and S.J. wrote the paper. S.J. supervised all aspects of the work.

## Declaration of generative AI and AI-assisted technologies in the manuscript preparation process

During the preparation of this work, the authors used ChatGPT in order to improve the grammar and readability. After using this tool, the authors reviewed and edited the content as needed and take full responsibility for the content of the published article.

