## Supplemental Material for "Protein Synthesis Profiling Enables In Vivo Capture of Ribosome-Associated mRNAs in Yeast"

**Table of Contents**

Supplementary Figures

- Supplementary Figure 1: RNA-seq normalization and sample-level quality control
- Supplementary Figure 2: DESeq2 interaction model analysis of AA and NOAA
- Supplementary Figure 3: Anota2seq model diagnostics and signal summaries
- Supplementary Figure 4: Metascape GO enrichment of translationally and transcriptionally regulated gene sets from PSP

**Supplementary Figures**

**Supplementary Figure 1: RNA-seq normalization and sample-level quality control.**

**(A)** VST mean–SD and expression density plots. The average Mean–SD (log–log) plots of VST-transformed expression values for total (TOT) and enriched (ENR) libraries in baseline control (BL), amino-acid–treated (AA), and NO amino-acid (NOAA) conditionsThe largely mean-independent spread of points indicates effective variance stabilization for downstream analysis.

**(B)** Density plot of VST-transformed gene expression values for each sample, grouped by condition (BL, AA, NOAA) and sequencing library type (total, T; enriched, E). Overlapping density curves. across all samples indicate successful normalization,absence of obvious technical biases or outlier libraries, and comparable expression distributions suitable for downstream linear modeling.

**(C)** Principal component analysis (PCA) of VST-transformed expression profiles. Each point represents a sample (BL, red circles; AA, green circles; NOAA, blue circles). Circle and triangle symbols indicate total and enriched libraries, respectively. Samples segregate primarily by treatment along PC1 (64% variance explained), with replicate 4 in each treatment displaced toward the edge of its cluster but not forming a separate cluster. This indicates that biological differences, rather than technical artifacts, drive the major variance components.

**(D)** Euclidean distance heatmap of inter- and intra-group sample relationships. Each cell represents the pairwise distance between VST-transformed expression profiles, with hierarchical clustering applied to samples. Replicates cluster tightly within conditions (BL, AA, NOAA) and sequencing types (TOT vs ENR) and are clearly separated between groups, confirming expected sample-to-sample similarity patterns and overall dataset integrity.

**Supplementary Figure 2: Condition-specific and shared interaction-model gene sets reveal distinct biological-process enrichment in AA and NOAA samples.**

**(A)** UpSet plot comparing significant genes from the AA and NOAA DESeq2 interaction-model analyses, separated by interaction-effect direction: AA_UP, AA_DWN, NOAA_UP, and NOAA_DWN. Vertical bars indicate unique or overlapping gene intersections, and horizontal bars indicate total genes per group. The largest intersections were unique to NOAA_UP and NOAA_DWN, indicating that the interaction-model results are dominated by condition-specific signatures, with fewer genes shared between AA and NOAA.

**(B)** Metascape Gene Ontology biological-process enrichment analysis of genes unique to each interaction-model gene set. Enriched terms are shown as a heatmap scaled by –log10(P). Unique NOAA_UP genes were enriched for protein localization, cell-cycle, kinetochore, and homeostatic processes, whereas unique AA_UP genes were enriched for organelle localization and protein-DNA complex assembly. Unique NOAA_DWN and AA_DWN genes were enriched for membrane targeting, ribosomal assembly, ribose phosphate metabolism, cytoplasmic translation, and translational fidelity. These results suggest that AA- and NOAA-specific interaction genes are associated with distinct biological programs.

**(C)** Metascape enrichment analysis of genes shared between AA and NOAA interaction-model datasets. Shared downregulated interaction genes were enriched for translation- and ribosome-associated processes, whereas shared upregulated interaction genes were associated with mRNA splicing, chromatin organization, and DNA-templated transcription. These shared signatures indicate common construct-dependent enrichment patterns associated with core gene-expression and RNA-processing pathways.

**Supplementary Figure 3: Anota2seq model diagnostics and signal summaries.**

**(A)** Density and frequency plots showing the distribution of raw and Benjamini–Hochberg (BH)–adjusted p-values for the omnibus interaction term in the anota2seq analysis of partial variance (APV) model. The raw p-values follow an approximately uniform distribution with no pronounced peak near zero, indicating that only a minority of genes exhibit interaction effects between treatment and total mRNA levels. The corresponding adjusted p-values cluster near one, supporting the common-slope assumption used by anota2seq.

**(B)** Density and frequency plots showing the distribution of raw and BH-adjusted Random Variance Model (RVM) p-values for the same omnibus interaction term. The approximately uniform distribution of raw RVM p-values suggests that variance moderation preserves the expected null behavior across genes. In contrast, the adjusted p-values cluster tightly near one, indicating that RVM shrinkage stabilizes variance estimates without inflating false positives.

**(C)** Random variance model (RVM) fits for interaction and omnibus group tests. For interaction terms (top) and omnibus group effects (bottom), Q–Q plots (left) compare empirical variance estimates with those expected under the inverse-gamma prior, and cumulative distribution plots (right) overlay empirical (black) and theoretical F-distributions (cyan) with Kolmogorov–Smirnov p-values (0.0191 and 0.0454, respectively). The close correspondence between empirical and theoretical curves indicates that the RVM provides an adequate description of variance across genes.

**(D)** Proportion of outliers in regression assessment using dfbetas. Bar plots show the observed (black) and simulated (red) proportions of data points exceeding several dfbeta cut-offs ($|dfb| > 1$, $> 2$, $> 3$, $> 2/\sqrt{N}$, $> 3/\sqrt{N}$, and $> 3.5 \times IQR$). The near-identical proportions indicate no excess of influential observations beyond that expected under a normal-error model, supporting the robustness of the APV regression fits.

**(E)** Density plots of raw p-values for total mRNA, translated mRNA, translational efficiency ("translation"), and buffering analyses in contrast 1. A prominent enrichment of very small p-values for total mRNA and a more modest enrichment for translated mRNA and translation indicate a strong transcript-level and moderate translational signal. In contrast, the flatter buffering distribution is consistent with fewer robust buffering events.

**(F)** Density plots of Benjamini–Hochberg FDR values for total mRNA, translated mRNA, translational efficiency, and buffering analyses in contrast 1. Strong enrichment of low FDRs for total and translated mRNA, a broader distribution for translation, and a peak near 1 for buffering indicate many transcript-level changes, fewer translational events, and relatively few high-confidence buffering events, consistent with the raw p-value distributions.

**(G)** Summary of all residuals from the APV regression model. Vertical bars show empirical residual quantiles across the standard normal distribution, with pink ticks indicating simulated normal envelopes. The expected proportion of outlier residuals is 1%, whereas 0.815% are observed, suggesting that residuals are approximately normally distributed and that the anota2seq model exhibits no excess of outliers.

**(H)** Residuals versus fitted values for the APV regression model. Each point represents a residual from the regression of translated on total mRNA as a function of its fitted value. Residuals form a roughly horizontal band centered around zero with approximately constant spread across the fitted range, supporting assumptions of linearity and homoscedasticity of residuals in the anota2seq model.

**Supplementary Figure 4: Metascape GO enrichment of translationally and transcriptionally regulated gene sets from PSP.**

GO Biological Process enrichment heatmap for transcriptionally repressed (mRNA Abundance Down) and transcriptionally enhanced (mRNA Abundance Up) gene sets. Rows correspond to enriched biological processes and the two columns to the down- and up-regulated transcriptional gene lists; colors represent $-\log_{10}(P)$ values. This view highlights biological processes that are predominantly governed at the mRNA abundance level, allowing for direct comparison with translational enrichment patterns.

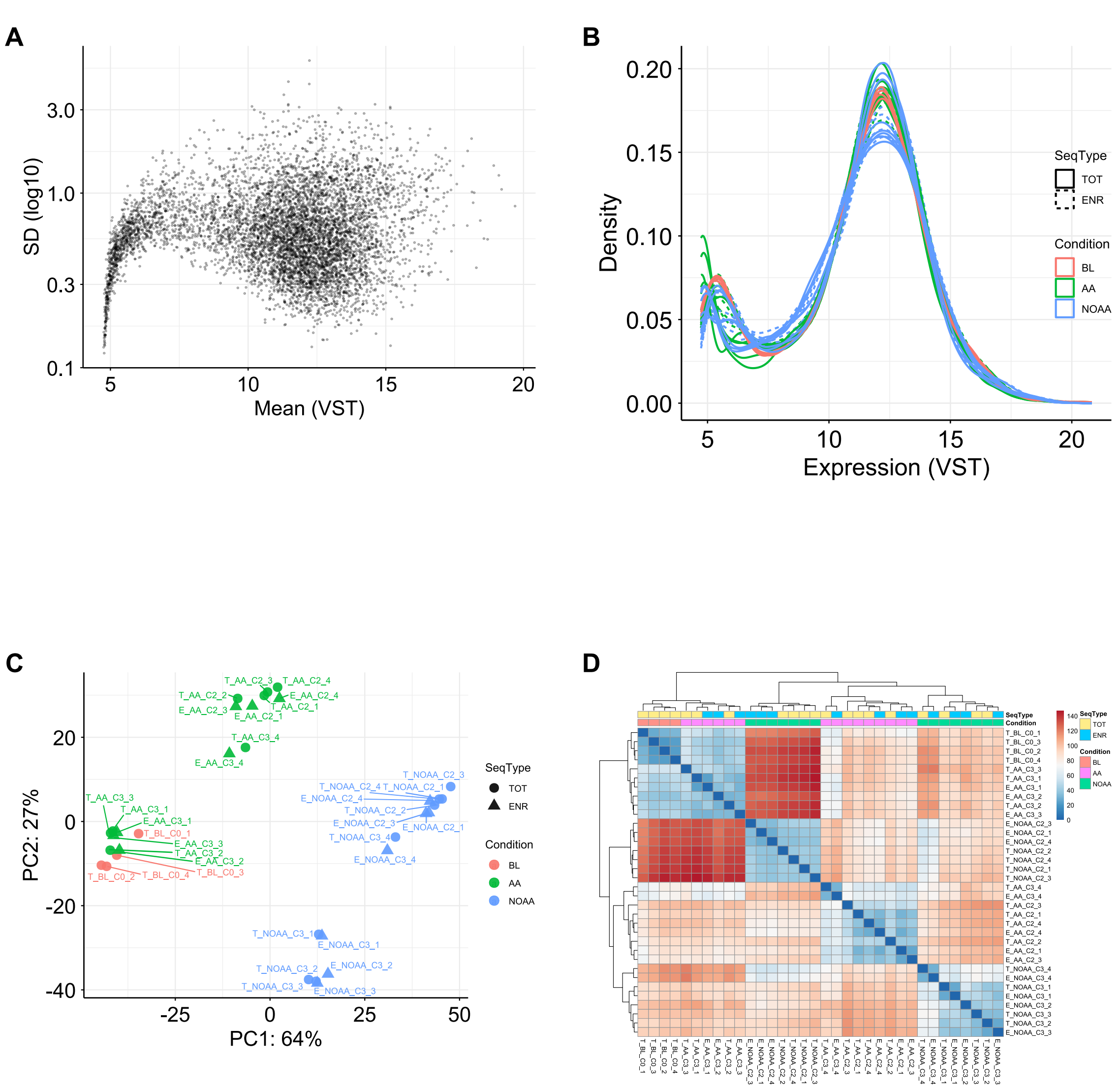


**Supplementary Figure 1**

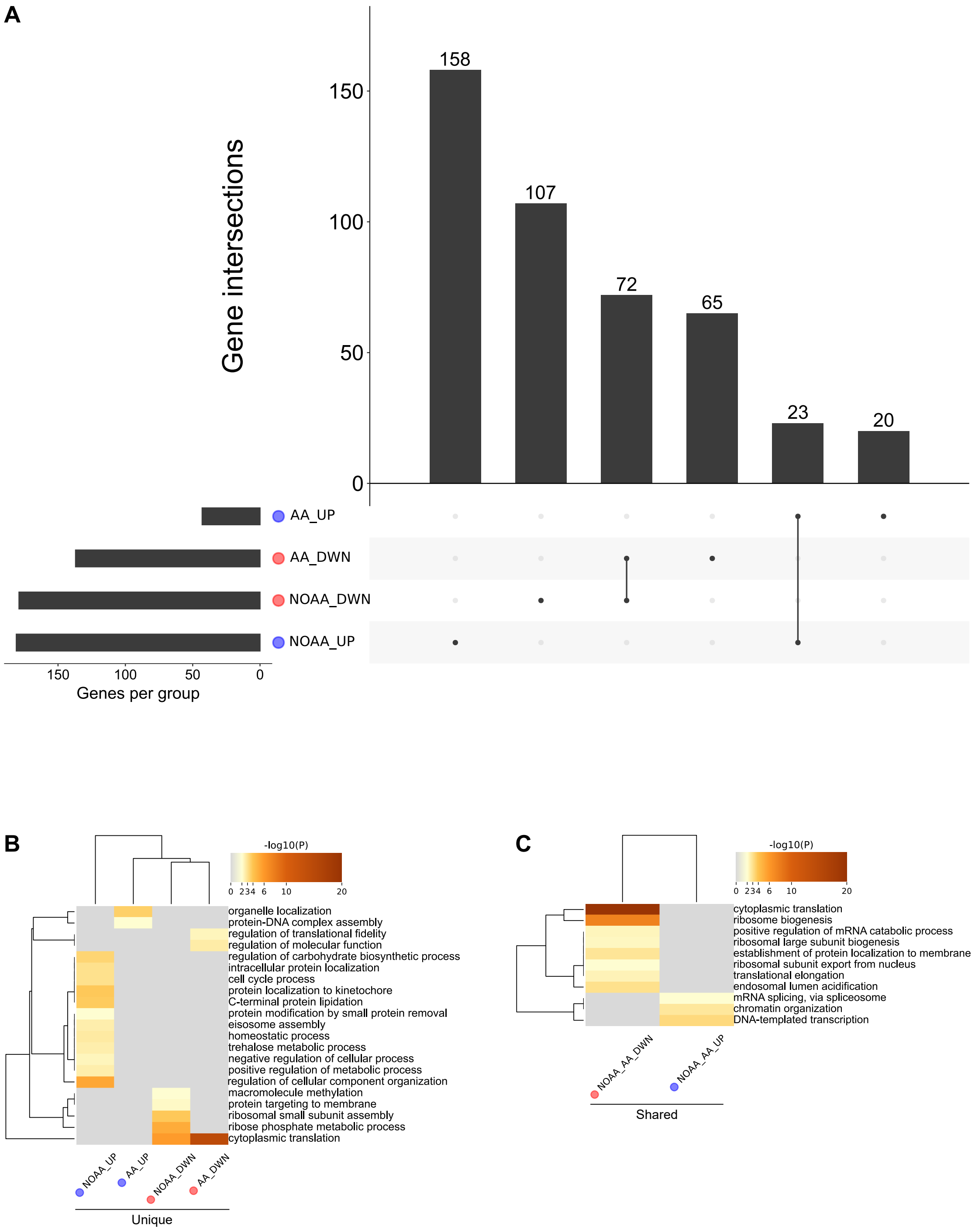


**Supplementary Figure 2**

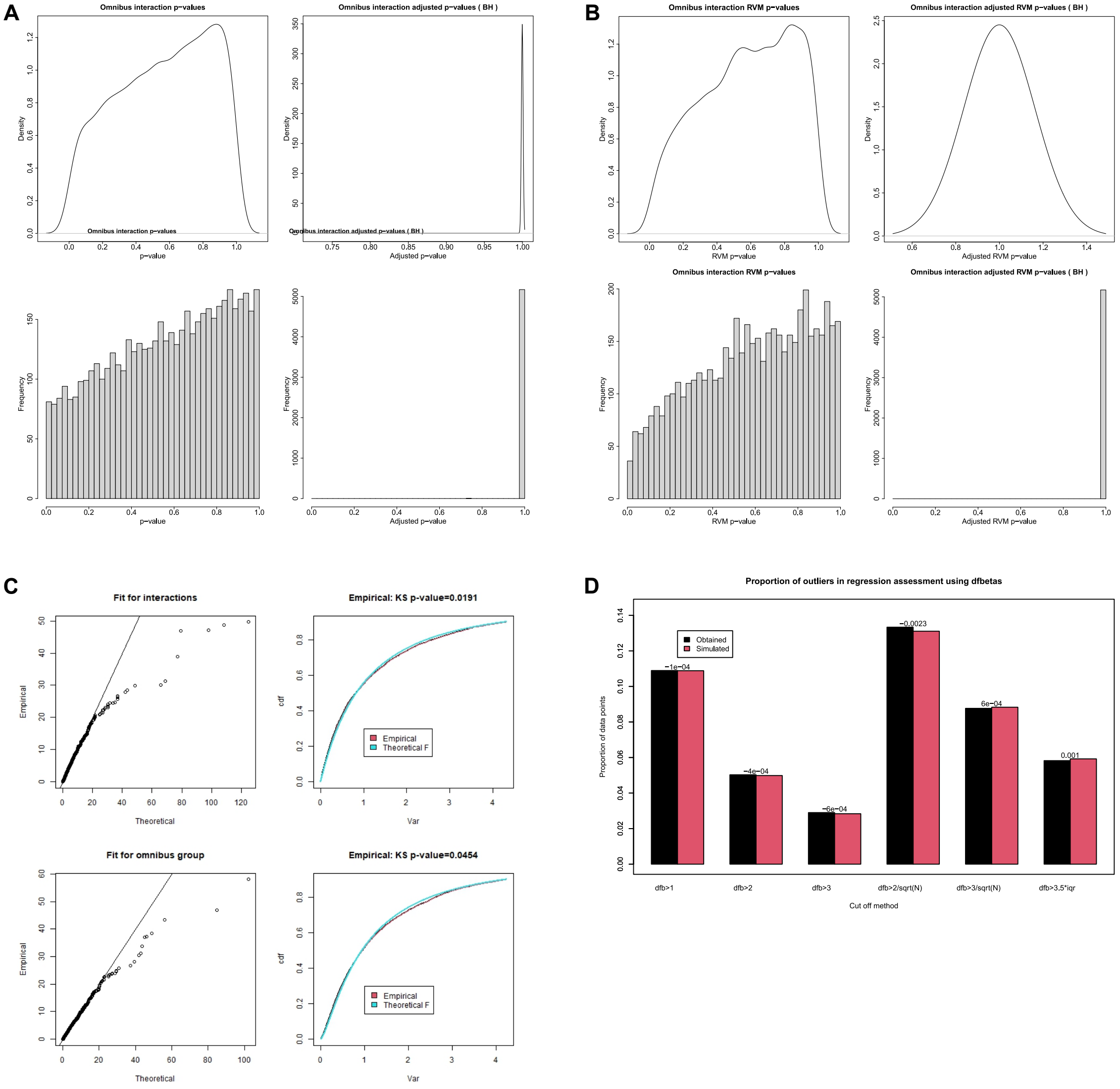


### Supplementary Figure 3

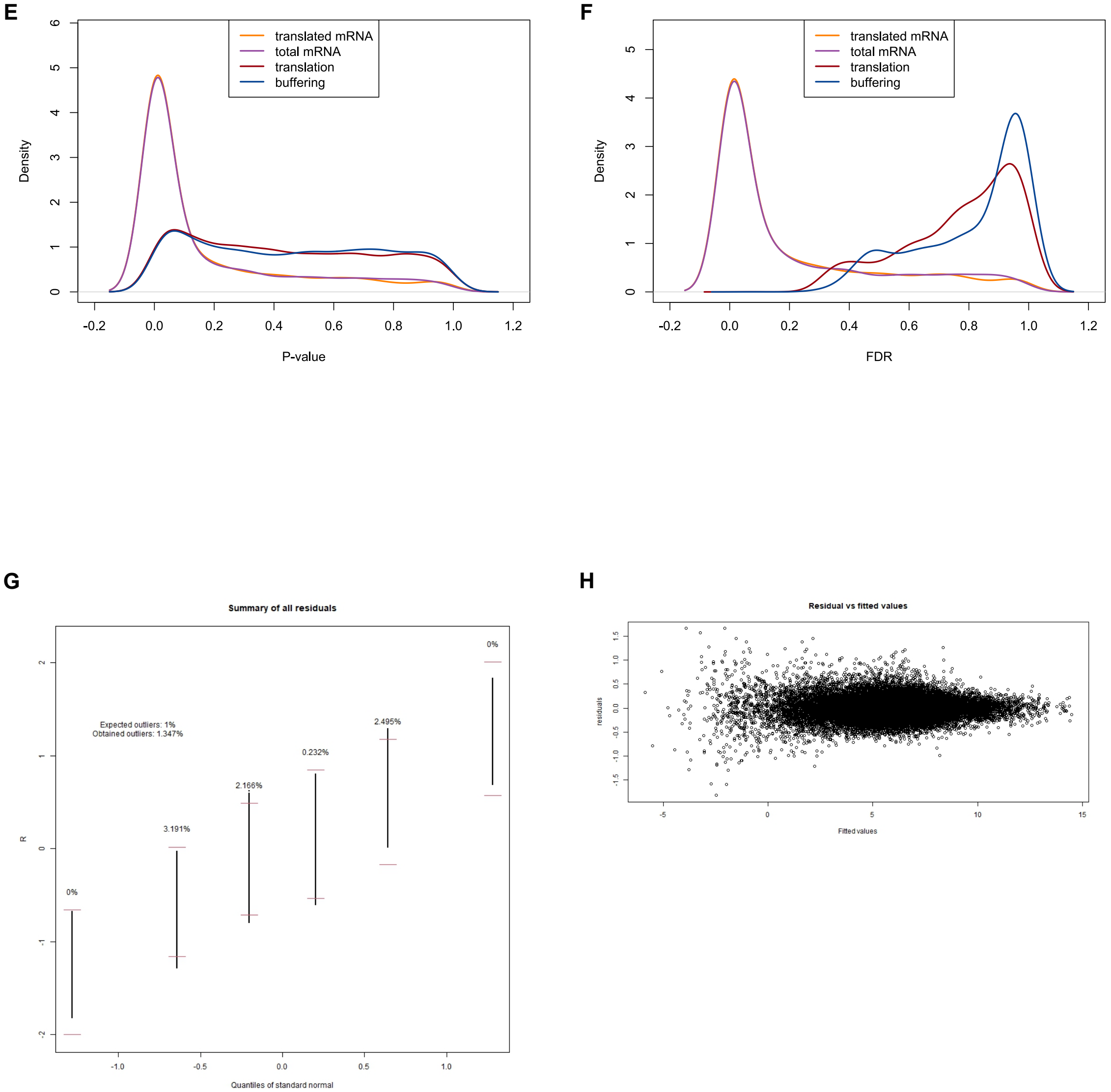


**Supplementary Figure 3 (Continued)**

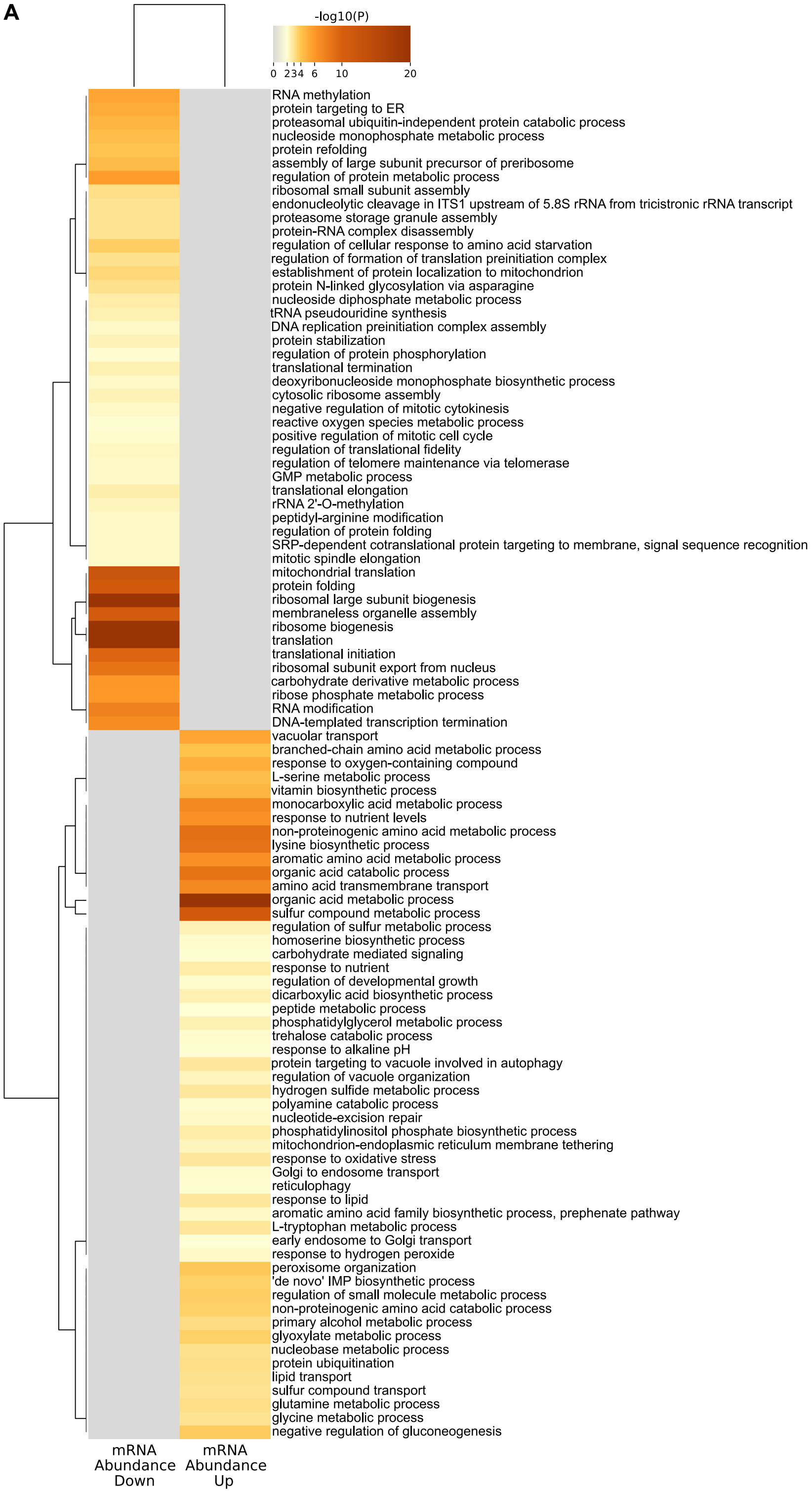


### Supplementary Figure 4
